# Structural insights into the ubiquitin independent MIDN-proteasome pathway

**DOI:** 10.1101/2025.03.04.641557

**Authors:** Nagesh Peddada, Xue Zhong, Yan Yin, Danielle Renee Lazaro, Jianhui Wang, Stephen Lyon, Jin Huk Choi, Xiaochen Bai, Eva Marie Y. Moresco, Bruce Beutler

## Abstract

The protein midnolin (MIDN) augments proteasome activity in lymphocytes, where it is predominantly expressed in adult mice. MIDN binds both to proteasomes and to substrates, but the mechanism by which MIDN facilitates substrate degradation in a ubiquitin-independent manner is unknown. Here, we present cryo-electron microscopy structures of the substrate-engaged, MIDN-bound human proteasome in two conformational states. MIDN induces proteasome conformations similarly to ubiquitinated substrates by using its ubiquitin-like domain to bind to the deubiquitinase RPN11. By simultaneously binding to RPN1 with its C-terminal α-helix, MIDN positions its substrate-carrying Catch domain above the proteasome channel through which substrates are translocated before degradation. Our findings suggest that both ubiquitin-like domain and C-terminal α-helix must bind to the proteasome for MIDN to stimulate proteasome activity.

## Introduction

The ubiquitin-proteasome system (UPS) is a regulated protein degradation system that removes damaged, misfolded, or otherwise unneeded proteins within cells. It functions during many cellular processes including cell division, differentiation, gene expression regulation, and proteotoxic stress management (1). The degradative component of the UPS is the proteasome holoenzyme, to which ubiquitinated and non-ubiquitinated substrates are recruited by ubiquitin (Ub) receptors on the proteasome, or by other mechanisms such as shuttling factors (2). The proteasome holoenzyme consists of the 20S proteolytic core particle (CP), and either one or two 19S regulatory particles (RP) capping the CP to form the 26S or 30S proteasome, respectively.

Extensive cryo-EM and crystal structural analyses have shown that the CP has a barrel shape formed by stacked rings of protein subunits (2–4). The top and bottom rings each consist of seven α subunits that form a gate closable to substrate entry. The two central rings each consist of seven β subunits, of which three possess proteolytic activity. The 19S RP contains 19 protein subunits including six AAA-family ATPase subunits (RPT1-6; aka PSMC1-6) and 13 non-ATPase subunits (RPN1-3, RPN5-13, and SEM1; aka PSMD subunits) (2, 5, 6). The ATPases form a hexameric ring positioned over the α-ring of the CP and serve to unfold and translocate substrate proteins through the center of the ring and into the CP (5, 6). Structurally, the ATPases assemble such that their oligonucleotide-binding sites form an upper-level ring (OB ring), while their AAA domains form a lower-level ring (AAA ring). The 13 RPN subunits form a hood-like structure extending from one side of the CP α-ring over the ATPase ring, with the exception of RPN1 that is bound to RPT1 in the ATPase ring opposite the hood (2, 7). The functions of the RPN proteins include acting as Ub receptors, deubiquitinases (DUB), and scaffold proteins.

Previous cryo-EM studies have captured the human proteasome in a variety of states that represent intermediates in the UPS cycle of substrate recruitment and degradation (3, 4, 8–13). These include substrate-free states (named S_A_, S_B_, S_C_, and S_D_) and substrate-engaged states (named E_A_, E_B_, E_C_, and E_D_), each of which consist of further sub-states in temporal sequence (e.g. E_A1_, E_A2_, etc.) (4, 8). Despite the abundance of structural data, dynamic regulation of protein degradation by the proteasome is incompletely understood, particularly the regulation of Ub-independent protein degradation (14). Recently, we and others showed that the protein midnolin (MIDN) interacts with the proteasome RP and enhances the proteolytic activity of the 26S proteasome (15, 16). MIDN also acts as a shuttling factor to recruit substrates to the proteasome in a Ub-independent manner. Immediate-early gene proteins such as EGR1 and FosB, and other transcription factors including IRF4 and NeuroD1, are substrates recruited by MIDN for proteasomal degradation (15). MIDN is abundant in lymphocytes and plays a crucial role in lymphocyte development and function (16). Moreover, MIDN may be an excellent therapeutic target in B cell malignancies because deletion of MIDN suppressed growth of MYC-driven B cell leukemia and a cultured myeloma cell line (16).

An AlphaFold-predicted structure of MIDN contains three folded domains: an N-terminal ubiquitin-like (UBL) domain, a C-terminal α-helix (αHelix-C), and a central folded domain containing several β-strands and α-helices previously named the Catch domain (Fig. S1, A and B) (15, 17). Biochemical assays demonstrated that the C-terminal α-helix was necessary and sufficient for MIDN binding to the proteasome, and all three structural domains were required to promote degradation of substrates by the proteasome (15, 16). However, no empirical structural data exist to explain the mechanism of substrate recruitment and proteasome activation by MIDN. In this study, we used cryo-EM to determine the atomic structure of MIDN-bound proteasomes. We present two proteasome conformational states upon MIDN binding, revealing that MIDN-bound proteasomes mimic the polyubiquitinated substrate engaged conformational state (E_B_ state) of the proteasome to promote substrate degradation.

### In vitro reconstitution and conformational states of MIDN bound proteasomes

Previously, we demonstrated that purified recombinant full-length human MIDN (468-aa isoform) enhances 26S proteasome activity by 3-4-fold in vitro (Fig. S1, C-E) (16). We confirmed that recombinant human MIDN directly interacts with purified human 26S proteasomes by pulldown assay (Fig. S1F). To capture substrate engaged human MIDN-proteasome complexes, we included purified recombinant human IRF4 as a substrate in our in vitro reconstitutions (Fig. S1G). MIDN-proteasome complexes were prepared following previously published protocols (8, 9) modified by the addition of a molar excess of a pre-mixed MIDN-IRF4 complex (Fig. S1H) to the 26S proteasome in the presence of ATPγS, with and without the 20S proteasome β subunit reversible peptide-like inhibitor MG-132. The complex was incubated for 5 min at room temperature and cryo-EM grids were plunge-frozen immediately. Additionally, as a control we prepared cryo-EM grids with the 26S proteasome in the presence of ATPγS and MG-132 but with neither MIDN nor substrate (MIDN-free proteasome, apo state). Overall, cryo-EM reconstructions and classifications for all datasets yielded seven proteasome states (Fig. S2A), which are further described below.

We collected an extensive dataset of MIDN- and substrate-bound proteasomes in the presence of ATPγS and MG-132, revealing predominantly two states (Fig. S3). The majority of particles (66%) corresponded to an S_B_-like state, while a smaller fraction (15%) represented an S_D_-like state. Further alignment-free 3D classification and focused local refinement (18) of the S_B_-like particles resulted in identification of three MIDN-bound E_B_-like states at nominal resolution ranging from 2.9 - 3.6 Å (Fig. S2A and S4A). Similarly, the S_D_-like particles were classified into three CP channel-open E_D_-like states at 3.8 - 4.2 Å resolution (Fig. S2A and S4A). We did not observe any E_A_-like particles in MIDN-proteasome dataset.

The cryo-EM densities of MIDN in our local resolution maps of the E_B_^MIDN^ states (Fig. S4A) were of adequate quality to construct atomic models of the MIDN C-terminal α-helix and UBL domain bound, respectively, to RPN1 and RPN11 (Fig. 1A-C). The MIDN-bound E_B_-like states (designated E_B1_^MIDN^, E_B2_^MIDN^, and E_B3_^MIDN^) represented substrate-engaged states of the 26S proteasome, as suggested by the formation of a quaternary subcomplex involving MIDN UBL domain-bound RPN11, RPN8, and the N-loop of RPT5 (Fig. 1D) (8). In all E_B_^MIDN^ states, we detected unfolded polypeptide density within the AAA-ATPase channel (Fig. S5A). In addition, the C-terminal tails (C-tails) of only three ATPases (RPT2, RPT3, and RPT5) were inserted into the inter-subunit pockets in the α-ring (α-pockets) of the CP, consistent with the closed CP gate (Fig. S5B). Since the three MIDN-bound E_B_-like states were distinguished by only subtle conformational differences (Fig. S2, B and C), hereafter we refer generically to an E_B_^MIDN^ state; in Figures we depict E_B1_^MIDN^ for E_B_^MIDN^.

**Figure 1.**
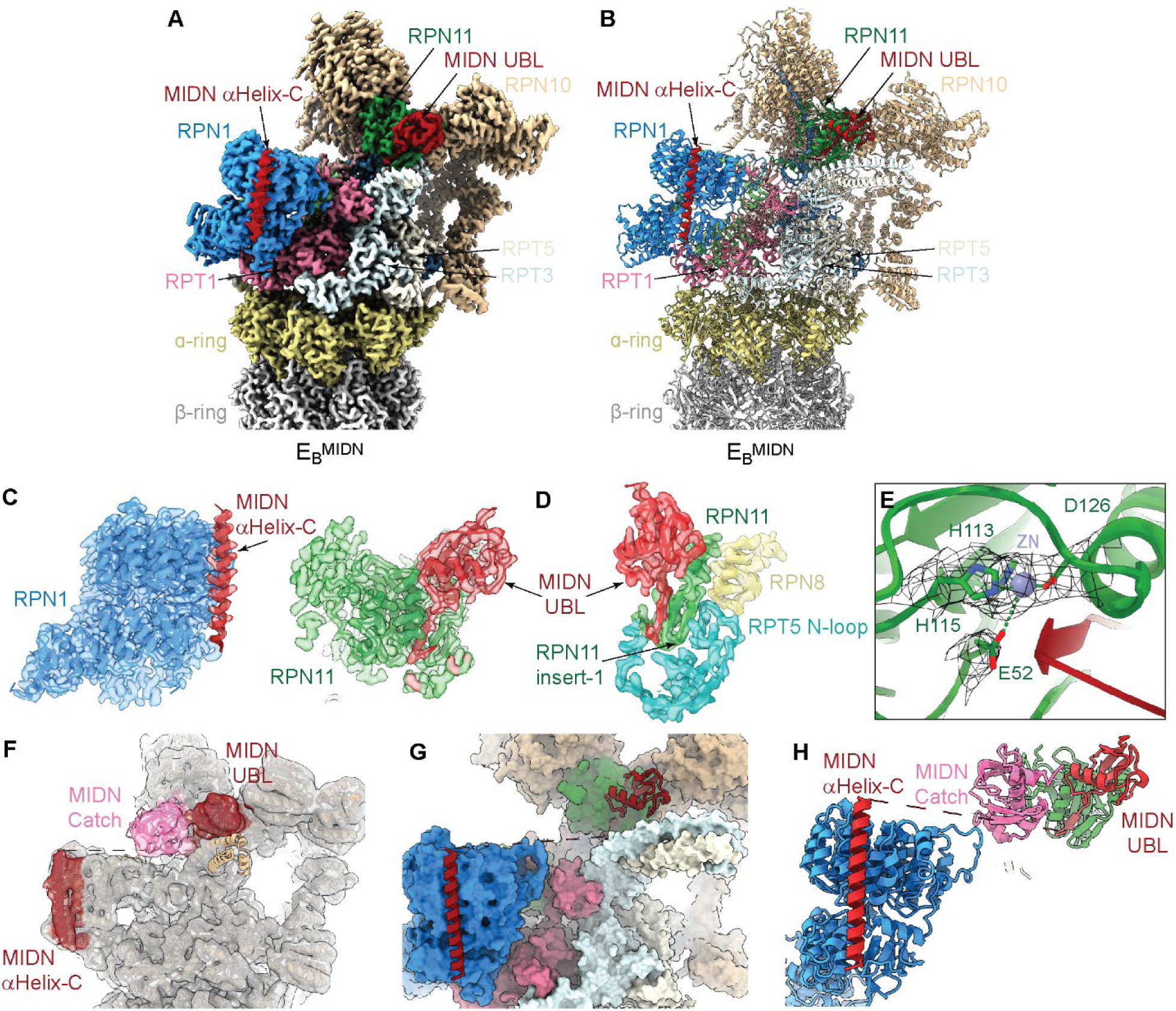
Cryo-EM reconstruction of substrate engaged MIDN-bound 26S proteasome complex. (A and B) Cryo-EM map (A) and atomic model (B) of substrate engaged MIDN-bound human proteasome complex in state E_B_^MIDN^. (C) Locally refined cryo-EM map is shown as a transparent surface overlayed with the cartoon representation of the atomic model of (left) RPN1 (dodger blue)-MIDN αHelix-C (brick red), and (right) RPN11(forest green)-MIDN UBL domain (brick red) from the E_B_^MIDN^ state. (D) Zoom-in view of the quaternary interface between MIDN-UBL, RPN11, RPN8, and RPN5 in state E_B_^MIDN^. The cryo-EM density is rendered as a transparent surface superimposed with the cartoon representation of the atomic model. (E) A close-up view of the tetrahedral co-ordination of a zinc ion (purple sphere) with histidine and glutamic acid residues of RPN11. Side chains of RPN11 that coordinate with the zinc ion are labelled and densities are shown in grey mesh. (F) Cryo-EM map of state E_B_^MIDN^, low pass filtered to 6 Å to show electron density for MIDN Catch domain (pink), and docked with the AlphaFold-predicted Catch domain as a rigid body in the density. (G) Anchoring of MIDN onto the proteasome. Cartoon representation of MIDN αHelix-C and UBL domain (brick red) with surface model of the 26S proteasome highlighting interacting proteins RPN1 (dodger blue) and RPN11 (forest green). (H) Cartoon representation of MIDN αHelix-C (brick red), Catch domain (pink) and UBL domain (brick red).

We subclassified approximately 143,000 S_D_-like proteasome particles into three highly similar states designated E_D1_^MIDN^, E_D2_^MIDN^, and E_D3_^MIDN^ (Fig. S2D and Fig. S3). The E_D_^MIDN^ states (hereafter E_D1_^MIDN^ is depicted for E_D_^MIDN^) were distinguished by an open CP gate and the C-tails of five ATPase subunits (RPT1, RPT2, RPT3, RPT5, and RPT6) inserted into the α-pockets of the CP (Fig. S5B). The E_D_^MIDN^ state was identical to the previously reported E_D2_ proteasome state (6MSK), a state of processive substrate translocation (Fig. S6F) (8). Therefore, we rigid body fitted and refined the E_D2_ structure of the 26S proteasome in our E_D1_^MIDN^ cryo-EM density, and in addition, observed an extra electron density adjacent to RPN11 that we were unable to identify due to limited particle number and low resolution (Fig. S2, A and D and S4A). Based on our finding that the MIDN UBL domain bound to RPN11 in the E_B_^MIDN^ state, we speculate that this low-resolution electron density corresponds to the MIDN UBL domain.

In an analogous dataset of MIDN- and substrate-bound proteasomes with ATPγS but without MG-132, cryo-EM reconstruction and classification resulted in two classes of particles: S_B_-like particles (19%) and S_D_-like (38%) (Fig. S7). The S_B_-like state was refined to an E_B_ state that lacked the C-terminal α-helix density bound to RPN1 (and therefore designated E_B_^MIDN-UBL^) but was otherwise identical to the E_B_^MIDN^ proteasome complex in the presence of MG-132 (Fig. S6C). We observed that the majority of particles were in an S_D_-like state in the absence of MG-132 (Fig. S7C), in contrast to the majority of particles in the presence of MG-132 that were in an S_B_-like state (Fig. S3C). Despite the majority of particles being in an S_D_-like state without MG-132 (2-fold more than the number of E_B_^MIDN-UBL^ particles), the cryo-EM map of the RP was of low resolution, and we could not determine if MIDN was bound. Together, the findings from two datasets, one with and one without MG-132, suggest that MG-132 stalls the activity of MIDN-bound proteasomes and thereby permits us to capture more extensive interactions with MIDN. Moreover, the ability to obtain high resolution maps of MIDN bound to E_B_ state proteasomes but not E_D_ state proteasomes suggests that MIDN binds more stably to the proteasome prior to the substrate translocating state.

Finally, cryo-EM reconstruction and classification of the MIDN- and substrate-free 26S proteasome in the presence of ATPγS and MG-132 yielded two classes of apo particles (Fig. S8). An S_A_-like state (designated S_A_^Apo^; 56%) aligned well with the published substrate-free S_A_ state of the human proteasome (4) (5T0G) rather than with the substrate-bound E_A_ state (8) (6MSB) (Fig. S6, G and H). In the S_A_^Apo^ state, Rpn11 is positioned above the Rpt4–Rpt5 interface on the OB ring without blocking the substrate entry port. In addition, C-tails of only two ATPases (RPT3 and RPT5) are inserted into α-pockets of the CP (Fig. S5B). The second class of apo particles exhibited an S_D_-like state (designated S_D_^Apo^; 17%) (Fig. S8C).

Our structures of the CP in the S_A_^Apo^, E_B_^MIDN^, and E_D_^MIDN^ states are identical up to their measured resolution, but we observed structural changes in the β subunits of the CP when compared to the cryo-EM structures of closed-channel CP structures in E_B_ state (6MSE) and E_A_ state (7W39), likely because of the peptide-like inhibitor MG-132 in our samples. We were able to rigid body fit a structure of a human CP with MG-132 (8CVR) into the cryo-EM maps of the E_B_^MIDN^ and E_D_^MIDN^ states. We observed distinct density for MG-132, allowing us to place an atomic model into the density map. MG-132 bound to CP catalytic β-subunits (β1, β2, β5) in our cryo-reconstructions (Fig. S9), in agreement with previous studies (19).

### MIDN-proteasome interactions

In our structural studies, MIDN-bound proteasomes favored the E_B_ over the E_D_ state. The E_B_ state is described as a deubquitination-compatible, closed-CP channel state that exists just prior to initiation of substrate translocation (8). The E_D_ state is an open-CP channel substrate translocation state (8). This suggests that MIDN (and substrate) interactions with the proteasome, in the absence of Ub, induce the formation of a substrate-engaged proteasome state ready for substrate translocation (Fig. 1, B and C). In the E_B_^MIDN^ state, MIDN bridges RPN1 and RPN11, with the C-terminal α-helix and the UBL domain of MIDN attached to RPN1 and RPN11, respectively. High-resolution cryo-EM maps showed no density for the Catch domain, possibly due to its flexibility or adoption of many conformations. However, a blob like electron density near RPN11 and above the OB ring of the AAA-ATPases was observed in low-pass filtered maps (6 Å), enabling the manual docking of an AlphaFold-predicted model of the Catch domain (Fig. 1F-H). Consistent with a reported function in interacting with substrates, the position of the Catch domain over the OB ring could place substrates at the appropriate site for entry into the ATPase channel.

### C-terminal α-helix of MIDN binds to RPN1

The MIDN C-terminal α-helix (amino acid residues 381-409) was previously shown to be necessary and sufficient for MIDN to bind to the proteasome (15). The amino acid sequence of the C-terminal α-helix is highly conserved (Fig. S10A). We found that it formed an extensive binding interface with RPN1 and buried a solvent accessible area of approximately 1097 Å^2^ on RPN1 (Fig. 2, A and B). RPN1 binds via two sites (T1 and T2) within its toroidal domain to both Ub and UBL domains, and thereby acts as a scaffold for recruitment of substrates, shuttling factors, and deubiquitinases to the proteasome (8, 20). The T1 site binds to mono- or di-Ub moieties and to the UBL domain of Rad32. The T2 site (toroid helices H22, H23, H24) binds to the UBL domain of yeast deubiquitinating enzyme Ubp6 and its human homologue USP14 (9, 20). MIDN bound to the T2 site of RPN1, with Arg381, Lys391, Arg406, and Asn385 at the ends of the C-terminal α-helix forming charged interactions with RPN1; and Leu395, Leu399, and Val392 forming hydrophic interactions with RPN1 (Fig. 2, A and B). RPN1 residues Glu319, Leu426, Asp430, Ser435, Asp438, Lys441, Asp468, Leu465 and His472 mediated the interaction with MIDN.

**Figure 2.**
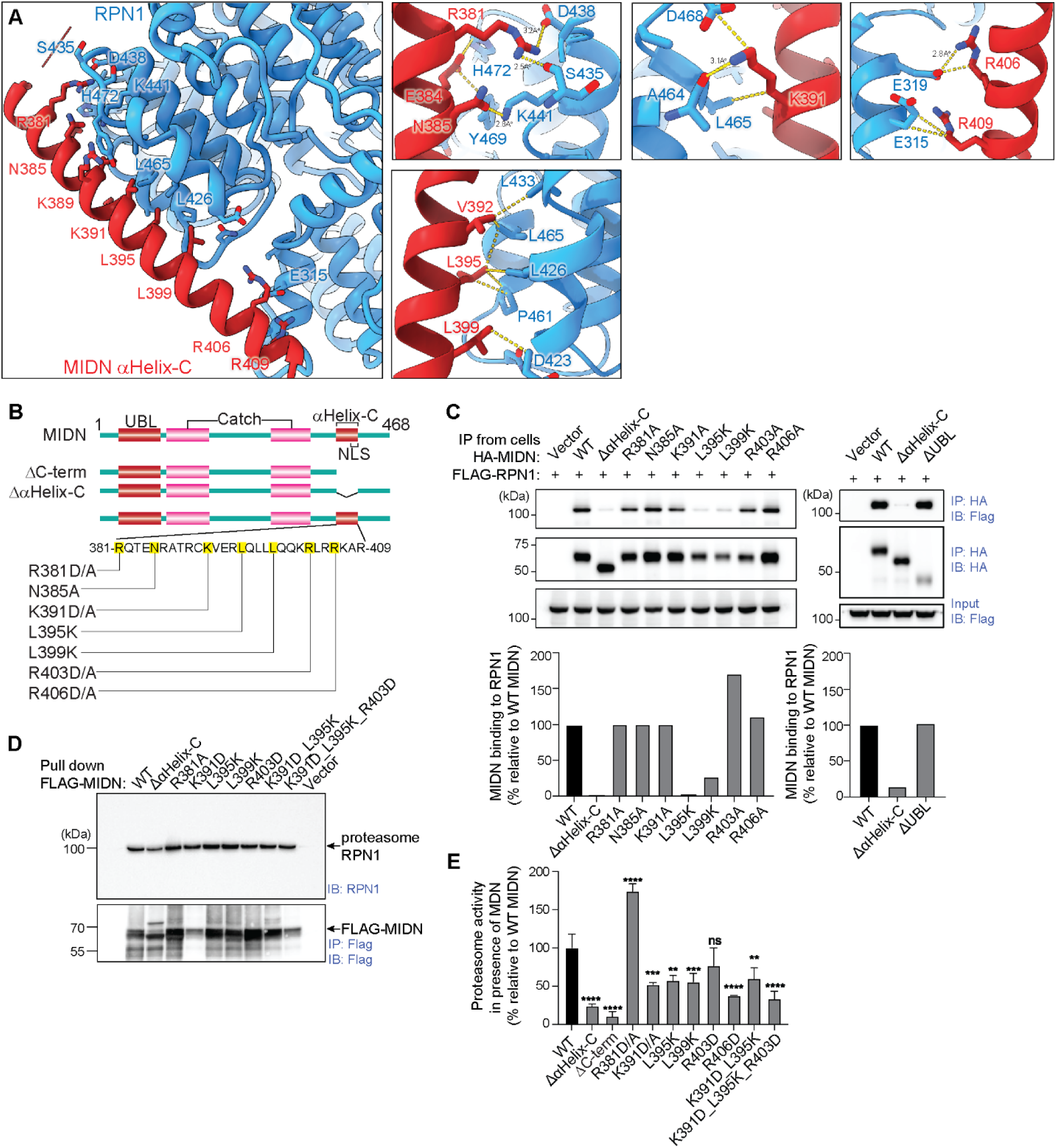
C-terminal *α*-helix of MIDN binds to RPN1. (A) Interface analysis of the interaction between MIDN αHelix-C and RPN1 in proteasome state E_B_^MIDN^. Side chains involved in interactions are labelled from both proteins. (Left) Overview of the interface. (Right, upper) Magnified views of individual amino acids Arg381and Asn385, Lys391, and Arg406 at the ends of the C-terminal α-helix forming charged interactions with RPN1. (Right, lower) Magnified view of Leu395, Leu399, and Val392 forming hydrophobic interactions with RPN1. RPN1 residues Glu319, Leu426, Asp430, Ser435, Asp438, Lys441, Asp468, Leu465 and His472 mediated the interaction with MIDN. (B) Schematic representation of domain organization of MIDN and amino acid sequence of MIDN αHelix-C. MIDN truncation mutants are depicted. The positions of point mutations tested in panels C-E are indicated in yellow. (C-D) Structure-based site directed mutagenesis experiments performed to test interaction of MIDN C-terminal α-helix mutants with RPN1. (C) (Upper) HEK293T cells transiently overexpressing HA-tagged WT or mutant MIDN, or Flag-tagged RPN1 were lysed, and then lysates were mixed and subjected to immunoprecipitation by anti-HA affinity gel and immunoblotted with anti-Flag or anti-HA. (Lower) Quantification of percentage of binding relative to WT MIDN was performed using ImageJ. (D) In vitro pull down of MIDN-proteasome complex. A 10-fold excess of purified 3X-Flag-tagged WT or mutant MIDN proteins were mixed with purified 26S proteasomes, immunoprecipitated with anti-Flag affinity gel, and immunoblotted with anti-RPN1 and anti-Flag. (E) In vitro peptidase activity of purified 26S proteasomes (1 nM) as measured by AMC fluorescence after hydrolysis of LLVY-AMC (10 μM) in the presence of ATP (0.2 mM). Purified WT or mutant MIDN was added at 100 nM, or no MIDN was added (Control). Activity is plotted relative to the activity of 26S proteasomes without added MIDN (Control, set at 100%) (n = 3 reactions per condition). P values were determined by one-way analysis of variance (ANOVA) with Dunnett’s multiple comparisons (E). ** P < 0.01; *** P < 0.001; **** P < 0.0001; ns, not significant.

We tested the importance of MIDN C-terminal α-helix residues for interaction with RPN1 by structure-based site directed mutagenesis (Fig. 2B). We confirmed that the C-terminal α-helix was necessary for MIDN to bind RPN1 (Fig. 2C). Point mutations L395K and L399K, affecting hydrophobic residues near the center of the helix, greatly diminished this interaction (Fig. 2C). In contrast, point mutations of positively charged residues (R381A, N385A, K391A, R403A, R406A) at either end of the helix individually did not reduce binding to RPN1 (Fig. 2C). We also tested the effect of these mutations on MIDN binding to the proteasome holoenzyme in vitro using purified proteins. As expected, we found that none of the mutations of the C-terminal α-helix abrogated MIDN binding to proteasome complexes, consistent with an ability of other MIDN domains to mediate binding (Fig. 2D). Finally, we tested whether MIDN with mutations of the C-terminal α-helix stimulated proteasome activity in vitro using the fluorogenic peptide substrate LLVY-AMC (Fig. S10D). Deletion of the C-terminal α-helix or the entire C-terminus resulted in a 70% reduction in the stimulatory effect of MIDN on proteasome activity compared to wild-type MIDN (Fig. 2E). Point mutations of the MIDN C-terminal α-helix, despite intact binding to the proteasome, stimulated proteasome activity at ∼50-60% of the WT MIDN level (Fig. 2E). The R381D/A mutation had no effect on binding to RPN1 or to proteasomes and enhanced MIDN stimulatory activity 1.75-fold (Fig. 2E). These data suggest that the C-terminal α-helix makes important contributions to the conformational regulation of proteasome activity by MIDN.

### UBL domain of MIDN binds to RPN11

RPN11 (PSMD14) is a proteasome resident deubiquitinase positioned close to the entrance of the ATPase channel in the proteasome RP. Previous work showed that substrate-conjugated Ub binds to RPN11 beginning in state E_A2_ and by state E_B_ has positioned its isopeptide bond within ∼3.5 Å of the RPN11 catalytic zinc ion, ready for cleavage (8). To achieve this, the insert-1 region of RPN11 forms a β-hairpin structure, which interacts with and aligns the C-terminus of Ub in the active site of RPN11. The MIDN UBL domain (amino acid residues 31-105) and Ub amino acid sequences do not share significant identity but adopt the same β-grasp fold (Fig. S10, B and C). We observed that the MIDN UBL domain bound to RPN11 in the same manner as ubiquitin in the E_B_ state of the substrate engaged 26S proteasome (Fig. 3, A and B) (8). These interactions are mediated by MIDN hydrophobic residues Thr38, Thr41, and Tyr43, and by the UBL domain C-terminal extension that projects down the RPN11 catalytic groove (Fig. 3A). An RPN11 α-helix (residues 103-137) and the β-hairpin loop formed by the insert-1 region interact with the UBL domain C-terminal extension, which takes the same position as the Ub C-terminus in the E_B_ state (Fig. 3B) (8). The UBL domain induces the same conformational changes in RPN11 and the ATPase ring as Ub itself does, and presumably positions the substrate for entry into the ATPase channel. Thus, the binding of the MIDN UBL domain to RPN11 appears to mimic ubiquitin-mediated substrate engagement, thereby allowing MIDN to facilitate ubiquitin-independent substrate degradation by the proteasome.

**Figure 3.**
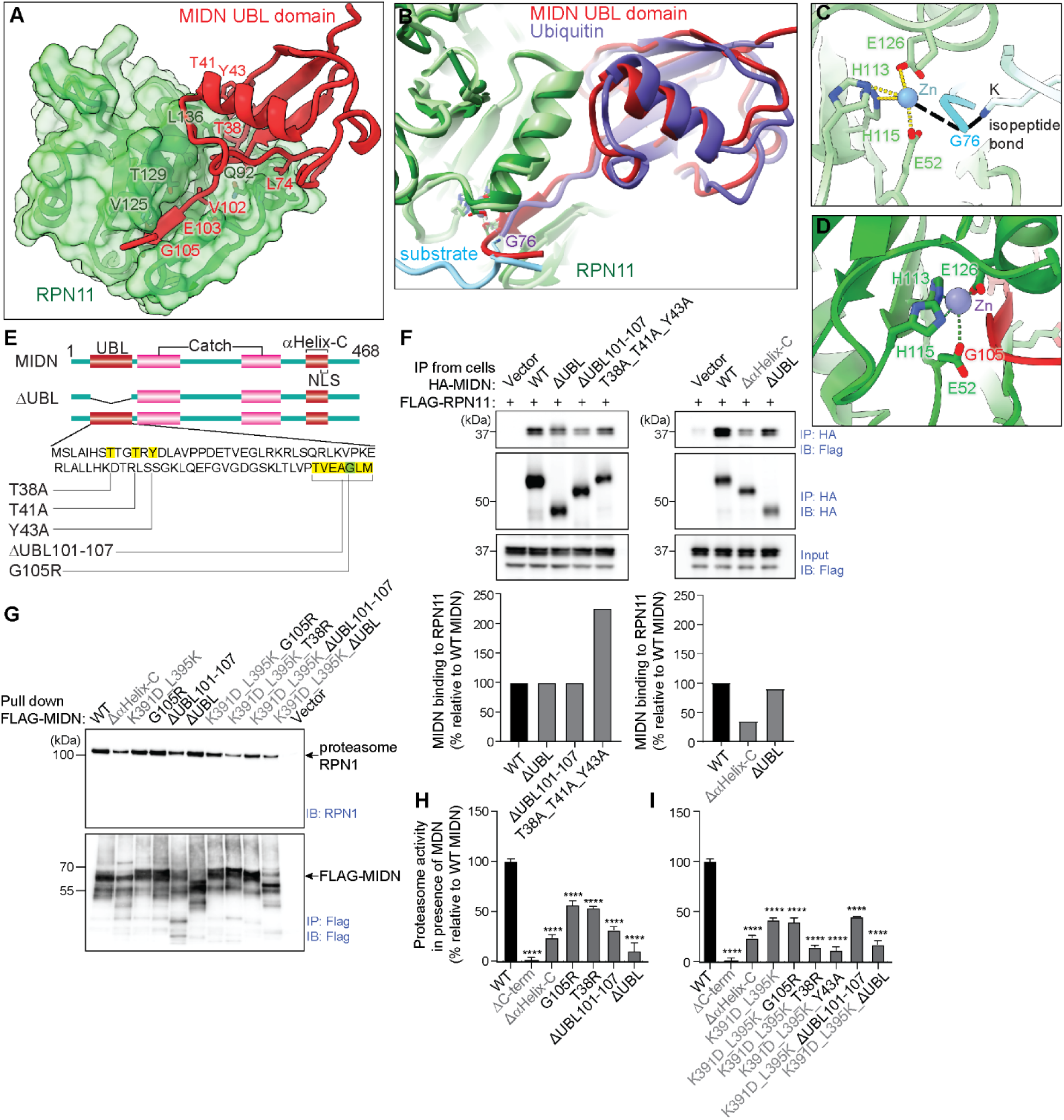
UBL domain of MIDN binds to RPN11. (A) Interface between MIDN UBL domain and RPN11 in proteasome state E_B_^MIDN^. Residues involved in the interaction between MIDN UBL domain and RPN11 are labelled (MIDN hydrophobic residues Thr38, Thr41, and Tyr43, and residues Val102, Glu103, and Gly105 of the UBL domain C-terminal extension that projects down the RPN11 catalytic groove). (B) Structural comparison of the MIDN UBL domain (brick red) bound to RPN11 (forest green) in proteasome state E_B_^MIDN^ with ubiquitin (purple) bound to RPN11 (light green) in proteasome state E_B_ (6MSE, substrate-engaged state). Gly76 of ubiquitin is labelled and substrate is shown in cyan from 6MSE structure. (C) Magnified view of catalytic site of RPN11 in substrate-engaged proteasome E_B_ state (6MSE). Gly76 of ubiquitin is involved in isopeptide bond formation and subjected to cleavage by RPN11, releasing the substrate from ubiquitin. (D) Magnified view of catalytic site of RPN11 in state E_B_^MIDN^, in which the corresponding glycine (Gly105) in the UBL domain of MIDN is not free to coordinate isopeptide bond formation with substrate and catalytic Zn ion. (E) Schematic representation of domain organization of MIDN and amino acid sequence of the MIDN UBL domain. MIDN ΔUBL truncation mutant is depicted. The positions of point mutations tested in panels F-I are indicated in yellow and green (G105). (F-G) Structure-based site directed mutagenesis experiments performed to test interaction of MIDN UBL domain mutants with RPN11. MIDN UBL domain mutations are labelled in black; MIDN αHelix-C mutations are labelled in grey. (F) (Upper) HEK293T cells transiently overexpressing HA-tagged WT or mutant MIDN, or Flag-tagged RPN11 were lysed, and then lysates were mixed and subjected to immunoprecipitation by anti-HA affinity gel and immunoblotted with anti-Flag or anti-HA. (Lower) Quantification of percentage of binding relative to WT MIDN was performed using ImageJ. (G) In vitro pull down of MIDN-proteasome complex. A 10-fold excess of purified 3X-Flag-tagged WT or mutant MIDN proteins were mixed with purified 26S proteasomes, immunoprecipitated with anti-Flag affinity gel, and immunoblotted with anti-RPN1 and anti-Flag. (H-I) In vitro peptidase activity of purified 26S proteasomes (1 nM) as measured by AMC fluorescence after hydrolysis of LLVY-AMC (10 μM) in the presence of ATP (0.2 mM). Purified WT or mutant MIDN was added at 100 nM, or no MIDN was added (Control). Activity is plotted relative to the activity of 26S proteasomes without added MIDN (Control, set at 100%) (n = 3 reactions per condition). (H) MIDN with either UBL domain or αHelix-C mutations. (I) MIDN with mutations of both αHelix-C and UBL domain. P values were determined by one-way analysis of variance (ANOVA) with Dunnett’s multiple comparisons (H, I). **** P < 0.0001.

Notably, point mutations, partial deletion, or full deletion of the UBL domain did not diminish binding of MIDN to RPN11 (Fig. 3, E and F), likely because the Catch domain also interacts with RPN11 (Fig. 1F). However, deletion of the C-terminal α-helix reduced binding to RPN11 by ∼60% (Fig. 3F); we speculate that the C-terminal α-helix may be required for initial docking of MIDN onto the proteasome which in turn enables the UBL domain to bind to RPN11. We also tested the binding of purified MIDN UBL domain mutants to the purified proteasome holoenzyme in vitro. As with mutations of the MIDN C-terminal α-helix, none of the tested UBL domain mutations abolished MIDN interaction with the proteasome (Fig. 3G). G76 in Ub is involved in isopeptide bond formation and subjected to cleavage by RPN11, releasing the substrate from Ub (Fig. 3C) (21). Mutation of the corresponding glycine (G105) in the UBL domain of MIDN (Fig. 3D) reduced the stimulatory effect of MIDN on proteasome activity by about 40%, whereas deletion of the UBL domain C-terminal extension (aa 101-107) reduced it by about 70% (Fig. 3H and Fig. S10D). A T38R mutation showed a similar effect as mutation of G105 (Fig. 3H and Fig. S10D). We also tested the effect of simultaneously mutating residues in the MIDN C-terminal α-helix (K391, L395) and the UBL domain, which consistently reduced the proteasome stimulatory activity of MIDN but in some cases not as severely as mutations of only one domain (Fig. 3I and Fig. S10D).

### Conformational dynamics of MIDN bound proteasome

Comparison of the MIDN-bound proteasome states with the S_A_^Apo^ state and with other previously reported proteasome structures showed that the RP conformations in the E_B_^MIDN^ and E_D_^MIDN^ states closely resemble those of the substrate-engaged MIDN-free E_B_ and E_D_ states, respectively (Fig. S6, B and F) (8). In contrast, compared to the S_A_^Apo^ state, in the E_B_^MIDN^ state the RP lid is rotated approximately 40° clockwise, resulting in rotation of the AAA ring with respect to the OB ring that in turn widens the ATPase axial channel and enhances substrate engagement (Fig. 4, A and B, and Fig. S6A). The USP14-proteasome structure is also distinct from the MIDN-proteasome structure (Fig. S6E), consistent with the idea that MIDN co-opts the RPN11-dependent pathway (mutually exclusive from the USP14 pathway) leading to substrate translocation (9). Our findings suggest that MIDN UBL domain binding emulates the binding of substrate-conjugated Ub to RPN11 and thus triggers the same changes in the ATPases that lead to substrate translocation.

**Figure 4.**
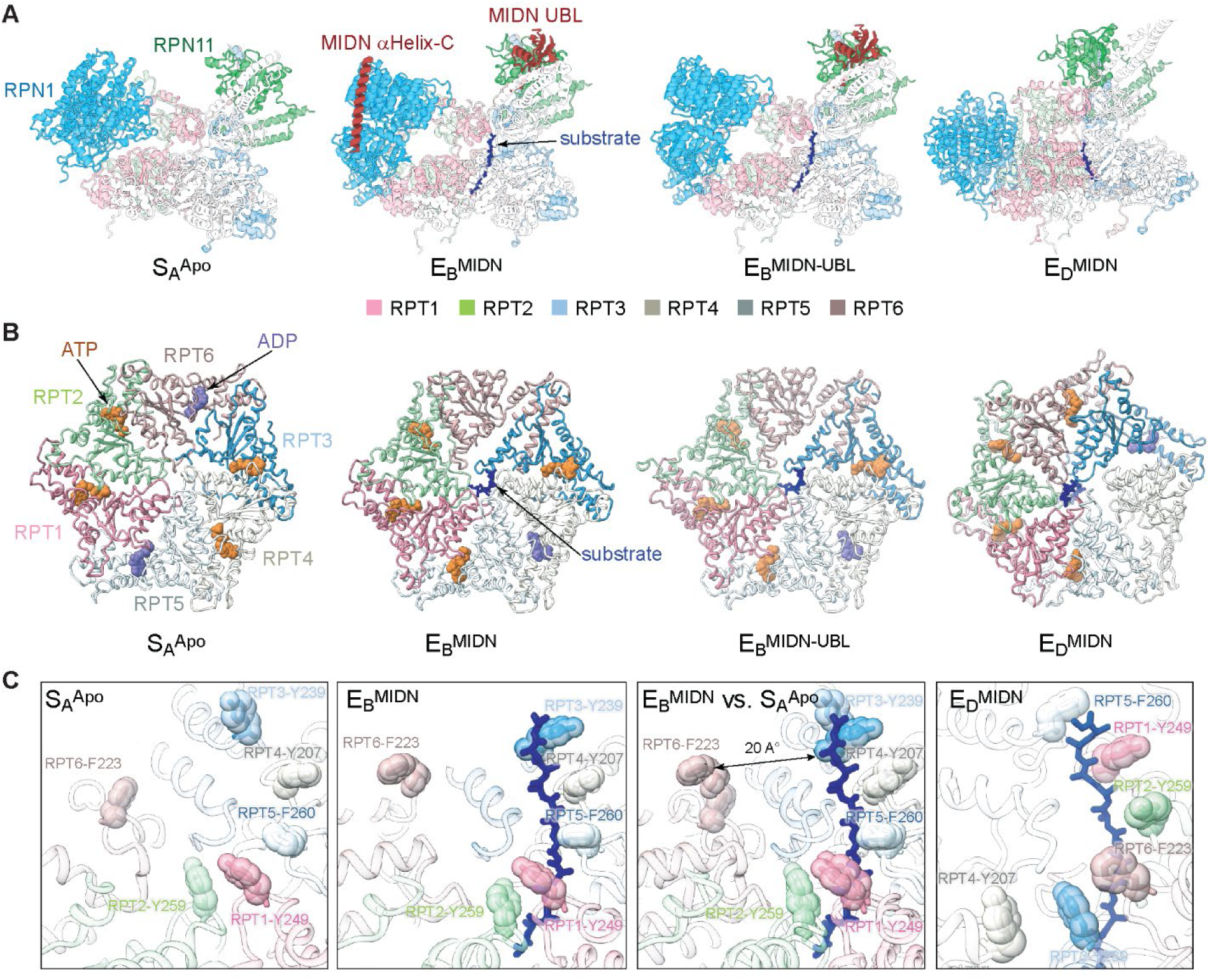
Conformational dynamics of MIDN bound proteasome. (A) Side by side comparison of the subcomplex containing MIDN, the six ATPases of the RP, RPN1, and RPN11 in different proteasome states. The CPs of the proteasome complex structures were superimposed and aligned, and the subcomplex orientations preserved in the images. The substrate is modelled as a polypeptide backbone and represented with blue sticks. (B) Top views of the AAA domain structures of the ATPases in distinct proteasome states. The substrate is shown in blue sticks. Nucleotides are shown in sphere representation (ADP, slate blue and ATP, yellow-orange). (C) Diverse configurations of the pore-1 loop staircase interacting with substrate in distinct proteasome states as compared to that in state S_A_^Apo^. Aromatic residues in the pore-1 loops are labelled and shown in stick representation superimposed with transparent sphere representation. The distance from a disengaged pore-1 loop to the substrate is marked. The third panel shows the superposition of states S_A_^Apo^ and E_B_^MIDN^.

Previous work showed that ATP hydrolysis by the ATPase subunits proceeds counterclockwise around the ring such that the ADP-bound state moves sequentially through each ATPase from RPT6 to RPT3 during a full cycle from state E_A_ to state E_D2_ (8). Our cryo-EM maps were of sufficient resolution to distinguish ADP from either ATP or ATPγS, but we could not discriminate the latter (Fig. S11). Similarly to the MIDN-free proteasome, we observed a counterclockwise rotation of the ADP-bound state of RPT proteins in the progression from S_A_^Apo^ → E_B_^MIDN^ → E_B_^MIDN-UBL^ → E_D_^MIDN^ (Fig. 4B). However, in E_B_^MIDN^ state we observed ATP bound to RPT2, which was different from the MIDN-free E_B_ state in which ADP was bound to RPT2 (Fig. 4B) (6MSE). The nucleotide states of the RPT subunits in our structures correlated with substrate binding and with the conformations of the pore loops that line the ATPase channel (Fig. 4C). In the S_A_^Apo^ state, the AAA channel has three narrow constrictions, defined by the axial area of overlap between the two pore-loop helices. In the E_B_^MIDN^ state, we observed substrate density in the AAA ring (Fig. S5A), the RPT subunits engaged with substrate are ATP-bound (Fig. 4, B and C), and an outward movement in the AAA domain widens the AAA channel (Fig. 4C). Comparison of E_B_^MIDN^ and E_D_^MIDN^ states shows that the pore-1 loop of RPT3 moves from the highest position to the lowest position in the substrate–pore loop staircase (Fig. 4C). These data support the idea that MIDN stimulates the canonical substrate-processing cycle of the proteasome, which is distinct from that of the proteasome allosterically activated by USP14 (9) or ZFAND5 (11).

### Model for the MIDN-proteasome pathway

Based upon our structural and functional data, we propose a model for MIDN-mediated, ubiquitin-independent proteasomal substrate degradation (Fig. 5). The sequence of proteasome states in this model is supported by their similarity to a previously defined cycle of ubiquitinated substrate processing by the 26S proteasome (4, 8). Moreover, the stepwise rotation of ATP hydrolysis by the AAA ATPases that we observed substantiates the proposed sequence of states in the model. First, a MIDN-substrate complex can bind to the proteasome S_A_^Apo^ state, which is a substrate-free closed-channel state (both ATPase and CP channels). The C-terminal α-helix and the UBL domain of MIDN form contacts with RPN1 and RPN11 subunits of the RP, respectively. Second, once bound to RPN11, the MIDN UBL domain induces the formation of the E_B_^MIDN^ state, a substrate translocation-competent state evidenced by the presence of substrate in the AAA domain channel.

**Figure 5.**
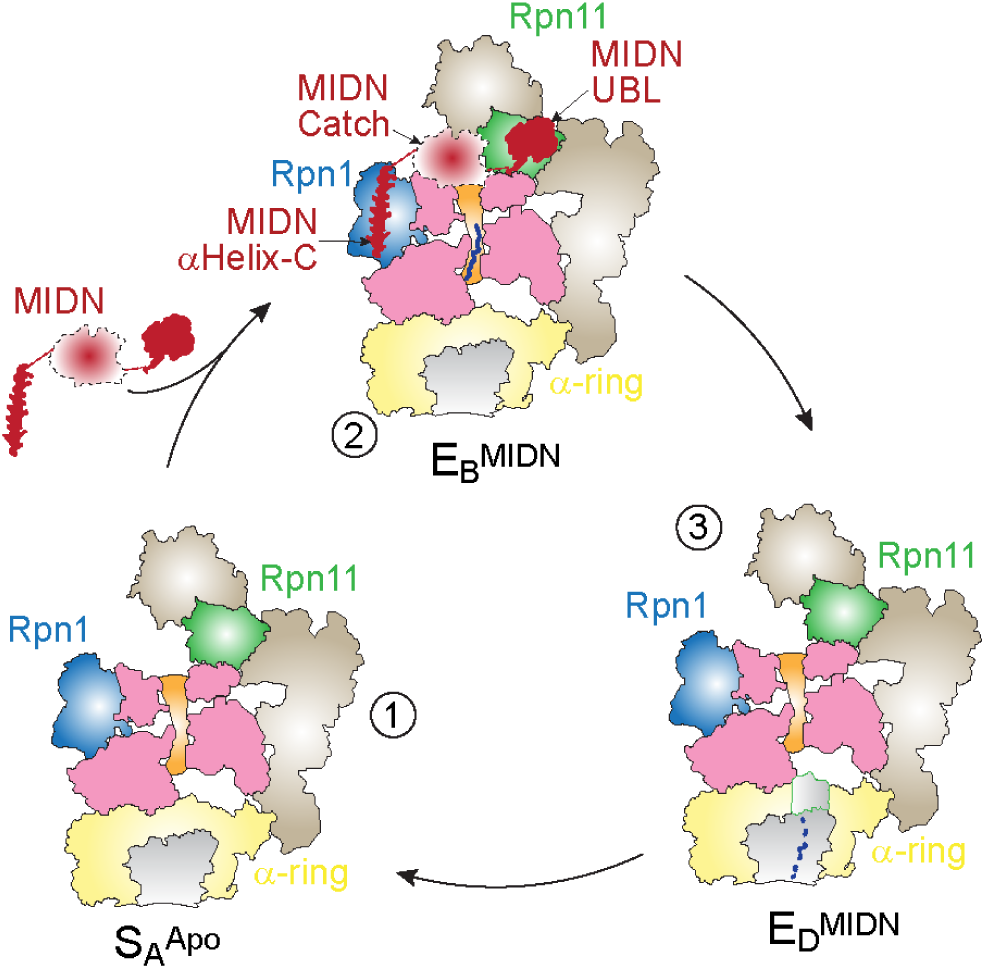
MIDN-mediated, ubiquitin-independent proteasomal substrate degradation. The sequence of proteasome states in the MIDN-mediated pathway is similar to those in a previously defined RPN11-catalysed ubiquitin-mediated 26S proteasome degradation pathway. MIDN promotes a catalytically active state of the proteasome (E_B_^MIDN^ state). First, a MIDN-substrate complex can bind to the proteasome S_A_^Apo^ state, which is a substrate-free closed-channel state. The C-terminal α-helix and the UBL domain of MIDN form contacts with RPN1 and RPN11 subunits of the RP, respectively. Second, once bound to RPN11, the MIDN UBL domain induces the formation of the E_B_^MIDN^ state, a substrate translocation-competent state evidenced by the presence of substrate in the AAA domain channel. The binding mode of the MIDN UBL domain mimics that of substrate-conjugated Ub to RPN11 and thereby enables MIDN to facilitate ubiquitin-independent substrate degradation while utilizing proteasome mechanisms that typically process ubiquitinated substrates. Third, the E_D_^MIDN^ state is presumably a substrate translocating state in which MIDN is motile by comparison to the E_B_^MIDN^ state.

We did not detect any E_A_-like particles in our dataset, suggesting that MIDN directly induces an E_B_ state. In contrast, other UBL domain-containing proteins, such as USP14/Ubp6 and RAD23, use their UBL domains to bind to RPN1. In the case of USP14, this binding allows the USP14 USP domain to interact dynamically with the AAA domain of RPT1, allosterically activating ATPase activity (9). The binding mode of the MIDN UBL domain mimics that of substrate-conjugated Ub to RPN11 and thereby enables MIDN to facilitate ubiquitin-independent substrate degradation while utilizing proteasome mechanisms that typically process ubiquitinated substrates. Third, the E_D_ ^MIDN^ state is presumably a substrate translocating state in which MIDN is motile by comparison to the E_B_^MIDN^ state. We speculate that MIDN is dislodged from the substrate during substrate translocation and dissociates from the proteasome. Importantly, we and others have demonstrated that all three domains of MIDN (UBL, Catch, and αHelix-C) are necessary for MIDN to stimulate proteasome activity and substrate degradation (15, 16). Our structural and mutagenesis data suggest this may be because the binding conformations of the C-terminal α-helix to RPN1 and the UBL domain to RPN11 are both important to induce ATPase and/or CP channel opening and to position the substrate-carrying Catch domain directly above the RP ATPase channel. Future structural and biochemical studies will be needed to test these hypotheses.

## Materials and Methods

### Plasmids

All constructs used here were created by standard molecular biology procedures, verified by sequencing to confirm the absence of undesirable mutations. Constructs used for recombinant protein production are described in detail in Table S1. Briefly, full length human MIDN (1-468 aa) and full length human interferon regulatory factor 4 (IRF4, 1-451 aa) cDNA sequences were synthesized (IDT) and used as templates to clone 3X-Flag tagged full length human MIDN^WT^, MIDN point mutants, and truncated versions of MIDN, and full length IRF4 into a customized pET-derived vector for bacterial overexpression. The vector contained an overexpression cassette encoding an N-terminal 14X-histidine followed by bdNEDD8 protease cleavage site. Point mutations and truncated versions were generated by standard site-directed mutagenesis using the Q5 Site-Directed Mutagenesis Kit (New England Biolabs).

### Expression and purification of recombinant MIDN protein and its variants

Purification of recombinant proteins was performed as described previously with slight modifications (16, 22). Customized pET-derived vectors encoding full length human MIDN^WT^, MIDN point mutants, and truncated versions of MIDN were transformed into *E. coli* BL21 Rosette (DE3) cells. Cells were grown at 37°C to an OD_600_ of 0.8 and induced with a final concentration of 0.25 mM IPTG (isopropyl-β-D-thiogalactoside) at 18°C overnight. After harvesting cells by centrifugation for 20 minutes at 4,000g, cell pellets were resuspended in B1 buffer (50 mM Tris pH 7.5, 800 mM NaCl, 40 mM imidazole, 10% glycerol) with complete protease inhibitors (Thermo Scientific™ Pierce Protease Inhibitor Tablets) and lysed via sonication. The lysate was cleared by centrifugation for 40 minutes at 18,000g. The cleared lysate was loaded onto a 5 mL Hi-Trap Ni-NTA column (Cytiva), washed with 5 column volumes of B1 buffer, and subsequently eluted with 2 column volumes of B2 buffer (50 mM Tris pH 7.5, 100 mM NaCl, 400 mM imidazole, 10% glycerol). The eluate was loaded directly onto a 5 mL Hi-Trap SP column (Cytiva). Protein was eluted using a linear gradient of buffer A1 (50 mM Tris pH 7.5, 50 mM NaCl, 40 mM imidazole, 10% glycerol) and buffer B1 spanning 10 column volumes. The main peak eluted from ion exchange was pooled, and affinity/solubility tags were cleaved overnight with bdNEDD8p protease. The proteins were purified by size exclusion chromatography on an S200 16/60 column equilibrated in gel filtration buffer (30 mM Tris pH 7.5, 150 mM NaCl, 1 mM TCEP). Purity was assessed by Coomassie stained SDS-PAGE. Aliquots were concentrated using Amicon Ultra spin columns (Merck Millipore) with a 30 kDa cutoff and snap frozen in liquid nitrogen for further use.

### Expression and purification of recombinant IRF4 protein

Expression and purification of recombinant full length human IRF4 were performed as described above.

### Cell culture, transfection, co-immunoprecipitation (co-IP), and immunoblotting

Cell culture, co-IP, and immunoblotting were described previously (23): HEK293T cells were grown at 37°C in DMEM (Life Technologies)/10% (v/v) FBS (Gibco)/1% antibiotics (Life Technologies) in 5% CO_2_. Transfection of plasmids was carried out using Lipofectamine 2000 (Life Technologies) according to the manufacturer’s instructions. 36 to 48 h after transfection, cells were harvested in NP-40 lysis buffer (20 mM Tris–Cl pH 7.5, 150 mM NaCl, 1 mM EDTA, 1 mM EGTA, 1% (v/v) Nonidet P-40, 2.5 mM Na_4_P_2_O_7_, 1 mM C_3_H_9_O_6_P, 1 mM Na_3_VO_4_, and protease inhibitors) for 45 min at 4°C. Co-IP assays were performed using cell extracts from HEK293T cells overexpressing RPN1, RPN11, MIDN^WT^, MIDN^△α-Helix-C^, MIDN^R381A^, MIDN^N385A^, _MIDNK391A, MIDNL395K, MIDNL399K, MIDNR403A, MIDNR406A, MIDN△UBL, MIDN△UBL101-107, or_ MIDN^T38A_T41A_Y43A^ proteins in separate cultures. Extracts of cells expressing the proteins of interest were mixed. Proteins were immunoprecipitated by anti-HA affinity gel (Sigma Aldrich). Captured proteins were eluted in PBS with 30 μL 200 μg/mL 2X-HA peptides (Sigma Aldrich), mixed with 8 μL of sample buffer with 0.1% bromophenol blue, heated to 95°C for 3 min, centrifuged at 12,000g for 1 min, and the supernatants were separated by SDS-PAGE for immunoblot analysis using anti-Flag (Sigma Aldrich, clone M2) or anti-HA (Cell Signaling Technology, clone C29F4) antibody. The membrane was incubated with secondary antibody at 1:4000 dilution [goat anti-mouse IgG-HRP (Southern Biotech) or goat anti-rabbit IgG-HRP (Thermo Fisher Scientific)] for 1 h at room temperature. The chemiluminescence signal was developed using SuperSignal West Dura Extended Duration Substrate kit (Thermo Fisher Scientific) and detected by a G:Box Chemi XX6 system (Syngene).

### MIDN-IRF4 pulldown

Purified 3X-Flag-MIDN^WT^ (5 μM) was mixed with purified IRF4 (10 μM) at a 1:2 ratio in buffer containing 50 mM Tris-HCl pH 7.5, 100 mM KCl, 1 mM TECP, and incubated for 1 h at 4°C with rotation. The mixture was passed through anti-Flag affinity gel (Sigma) and bound complex was eluted using buffer containing 50 mM Tris-HCl pH 7.5, 100 mM NaCl, 1 mM TECP, and 150 μg/mL 3X-Flag peptide. Eluted fractions were assessed by Coomassie stained SDS-PAGE and immunoblotted with anti-IRF4 antibody (Cell Signaling Technology, clone D9P5H) and anti-Flag antibody (Sigma Aldrich, clone M2).

### Proteasome stimulating activity assay

Proteasome stimulating activity of MIDN towards human proteasomes (26S) was measured using fluorogenic substrate Suc-LLVY-AMC (South Bay Bio) as described previously (16). Briefly, 1 nM human 26S proteasomes (Ubiquitin-Proteasome Biotechnologies) were incubated with 100 nM of 3X-Flag tagged full length human MIDN^WT^, MIDN point mutants, or truncated versions of MIDN (Table S1) in buffer (50 mM Tris-HCl pH 7.5, 100 mM KCl, 0.5 mM MgCl_2_, 1 mM TECP, 0.2 mM ATP and 25 ng/μL BSA) for 5 min at room temperature. 10 μM Suc-LLVY-AMC (South Bay Bio) was added to the reaction mixture. For time course analyses, proteasome activity was measured beginning immediately after substrate addition and for up to 1.5 h at 37°C using a BioTek multimode plate reader (excitation, 345 nm; emission, 445 nm).

### Pulldown of MIDN-26S proteasome complex

A 10-fold excess of purified 3X-Flag tagged full length human MIDN^WT^, MIDN point mutants, or truncated versions of MIDN was mixed with purified 26S proteasomes (South Bay Bio) in buffer containing 50 mM Tris-HCl pH 7.5, 100 mM KCl, 5 mM MgCl_2_, 1 mM TECP, 10% glycerol, and 1.5 mM ATPγS and incubated for 1 h at 4°C. Each mixture was passed through anti-Flag affinity gel (Sigma) and bound complex was eluted using buffer containing 50 mM Tris-HCl pH 7.5, 100 mM KCl, 5 mM MgCl_2_, 1 mM TECP, 1.5 mM ATPγS, and 150 μg/mL 3X-Flag peptide. Eluted fractions were assessed by Coomassie stained SDS-PAGE and by immunoblot analysis using anti-Flag (Sigma Aldrich, clone M2) or anti-RPN1 (Cell Signaling Technology, clone 14141) antibody. The membrane was incubated with secondary antibody at 1:4000 dilution [anti-mouse IgG-HRP (Sigma) or anti-rabbit IgG-HRP (Thermo Fisher Scientific)] for 1 h at room temperature. The chemiluminescence signal was developed using Super Signal West Dura Extended Duration Substrate kit (Thermo Fisher Scientific).

### Cryo-EM sample and grid preparation

To prepare MIDN-proteasome complexes for cryo-EM, human 26S proteasomes (Ubiquitin-Proteasome Biotechnologies, purified from HEK293 cells) were buffer exchanged into 50 mM Tris pH 7.5, 150 mM NaCl, 20 mM KCl, 2 mM ATPγS, 5 mM MgCl_2_, and 1 mM TECP by using Zeba™ Micro Spin Desalting Columns (7K MWCO, Thermo Fisher 89883). All recombinant proteins were exchanged into 50 mM Tris pH. 7.5, 150 mM NaCl, and 1 mM TECP. The MIDN-IRF4 complex was made by mixing the proteins (7 μM) at a 1:1 ratio and incubating on ice. After buffer exchange, ∼50-fold molar excess of the MIDN-IRF4 complex was added to the 26S proteasomes followed by addition of the peptide-like inhibitor MG-132 (40 μM final concentration) to the mixture. After a 5-min incubation at room temperature, Nonidet P-40 (0.005% v/v) was added to the sample before plunge freezing the cryo-EM grids. For MIDN-IRF4-free 26S proteasomes, proteasomes were prepared as above except the addition of MIDN-IRF4 complexes was omitted. For MIDN-bound 26S proteasomes without MG-132, proteasomes were prepared as above except the addition of MG-132 was omitted.

Cryo-EM grids with MIDN-proteasome complexes in the presence of MG-132 or with MIDN-free proteasome complexes were prepared using 3.5 µL of sample (A_280_ of 1.66), applied to Quantifoil copper R1.2/1.3 300-mesh grids (Ted Pella) for cryo-EM data collection. Grids were glow discharged using a PELCO easiGlow unit (Ted Pella) at 30 mA for 80 s and blotted with the following parameters: blot force of -3 to 5, blot time 4.0 s. Grids were frozen in liquid ethane using a FEI Mark IV Vitrobot (Thermo Fisher) set at 4°C and 100% humidity. 3X-Flag enriched MIDN-proteasome complexes in the absence of MG-132 were prepared on continuous carbon Quantifoil copper 300 mesh (R 2/1) holey grids by applying a volume of 3-4μl of sample (A_280_ of 1.3) onto glow-discharged grids (30 mA for 30 s, blot force of 8-12, blot time 4.5 s, and wait time 15 s). Grids were plunged into liquid ethane using a FEI Mark IV Vitrobot (Thermo Fisher) at 100% humidity.

### Cryo-EM data collection and data processing

MIDN-bound and MIDN-free proteasome complex datasets were acquired on a FEI Titan Krios microscope (Thermo Fisher) operated at 300 kV and equipped with a post-column energy filter (Gatan) and a K3 direct detection camera (Gatan) in non-CDS mode using SerialEM (24). Movies were acquired at a pixel size of 0.54 Å in super-resolution counting mode, with an accumulated total dose of 50 e-/Å^2^ over 50 frames. The defocus range was set from −1.1 to −2.7 μm. Micrographs of MIDN-proteasome complexes without MG-132 were acquired on a FEI Titan Krios microscope (Thermo Fisher) operated at 300 kV with a Falcon 4 camera (Thermo Fisher Scientific), using a slit width of 10 eV. A calibrated magnification of 105,000 was used for imaging, yielding a pixel size of 0.738 Å on images. Each micrograph was dose-fractionated to a dose rate of 7.76 e-/pixel/s, with a total exposure time of 3.45 s, resulting in a total dose of about 50 e-/Å^2^. Movies were stored in EER format with 1092 raw frames per movie. The defocus range was set from −1.2 to −2.7 μm. 20,794 movies were collected for the MIDN-proteasome complex with MG-132 sample, 13,162 movies for the MIDN-free proteasome sample, and 17,856 movies for the MIDN-proteasome complex without MG-132 sample.

All data processing was performed using the software cryoSPARC v4 (25). To correct for beam induced motion and compensate for radiation damage over spatial frequencies, the patch motion correction algorithm was employed using a binning factor of 2, resulting in a pixel size of 1.08 Å/pixel for the micrographs. Contrast Transfer Function (CTF) parameters were estimated using patch CTF estimation.

### MIDN-26S Proteasome-ATPγS-MG-132 dataset

Data from three separate grids frozen at the same time (identical sample) were combined. A small subset of ∼20 frames was used to generate the initial template for particle picking. Initially, 3,102,828 particles were picked from 20,794 micrographs using template picker and extracted with box size of 640 pixels from all the micrographs; particles were binned four times and used for 2D classification. After two rounds of two-dimensional (2D) classification, around 939,289 particles were selected for subsequent *ab initio* modeling, followed by heterogeneous refinement (Fig. S3). After heterogenous 3D classification, we combined S_B_-like particles and S_D_-like single- and double-capped particles and re-extracted the particles with a pixel size of 1.08 Å for each class.

Individual classes were refined with cryoSPARC homogeneous refinement to a high resolution of 2.96 – 3 Å, followed by CTF refinement. To investigate the conformational states of the 19S regulatory particle (RP), we subtracted 20S core particle (CP) signal and performed focused refinement of the RP as described in previous studies (18). RP-focused alignment-free 3D classification of the RP of S_B_-like particles (623K) and S_D_-like particles (143K) yielded three E_B_-like states and three E_D_-like states, respectively, named in accordance with previous convention (8, 9). DeepEMhancer (26) was then used with the two unfiltered half maps to generate a locally sharpened map. To better resolve the interactions between MIDN and proteasome components, 232K E_B1_ particles were selected and masks were applied, centered on RPN1 and MIDN αHelix-C, and on RPN11 and MIDN UBL domain; signals outside the masks were subtracted (Fig. S3). Local refinement of these classes resulted in maps with overall resolutions of 2.8 Å, which were further sharpened by DeepEMhancer. These maps were then aligned with the full map and combined using the “vop maximum” function in UCSF Chimera based on the maximum value at each voxel (27). All resolutions were estimated by applying a soft mask around the protein density using the gold-standard Fourier shell correlation (FSC) = 0.143 criterion (28) (Fig. S4A).

### 26S Proteasome-ATPγS-MG-132 dataset

Initial 2D classification and processing were performed identically to MIDN-proteasome dataset. Initially, 1,092,420 particles were picked from 13,162 micrographs using template picker and extracted with box size of 640 pixels from all the micrographs; particles were binned four times and used for 2D classification. After two rounds of 2D classification, around 252,319 particles were selected for subsequent *ab initio* modeling, followed by heterogeneous refinement (Fig. S8). After heterogenous 3D classification, we combined single- or double-capped S_A_-like particles and S_D_-like particles and re-extracted the particles with a pixel size of 1.08 Å for each class. Individual classes were refined with cryoSPARC homogeneous refinement to a high resolution, followed by CTF refinement. RP-focused alignment-free 3D classification of the RP of S_A_-like particles (140K) yielded three S_A_-like states (two single- and one double-capped state), which were combined and further refined using homogenous refinement. S_D_-like particles (42,984) were not further classified due to their small number. To improve local resolution of the RP we subtracted CP signal from the particles and performed focused local refinement of the RP (18); the same procedure done for the CP. To better resolve the RPN1 density, 106K particles were selected, a mask was applied around the RPN1 density, and local refinement of RPN1 was performed. DeepEMhancer (26) was then used with the two unfiltered half maps to generate a locally sharpened map. These maps were then aligned with the full map and combined using the “vop maximum” function in UCSF Chimera based on the maximum value at each voxel (27) (Fig. S8). All resolutions were estimated by applying a soft mask around the protein density using the gold-standard Fourier shell correlation (FSC) = 0.143 criterion (28) (Fig. S4C).

### MIDN-26S Proteasome-ATPγS-dataset

Initial 2D classification and processing were performed identically to apo-state dataset. Initially, 1,574,488 particles were picked from 17,856 micrographs and extracted with box size of 700 pixels from all the micrographs; particles were binned four times and used for 2D classification. After the two rounds of 2D classification, around 471,439 particles were selected for subsequent *ab initio* modeling, followed by heterogeneous refinement (Fig. S7). After heterogenous 3D classification, we combined single- and double-capped S_B_-like particles (94K, 19%), and SD-like particles (178K, 38%) and re-extracted the particles with a pixel size of 0.86 Å for each class. Individual classes were refined with cryoSPARC homogeneous refinement to a high resolution of 2.6-2.9 Å, followed by CTF refinement. To investigate the conformational states of the RP, we subtracted CP signal from the particles and performed focused refinement of the RP as described in previous studies (18); the same procedure done for the CP. DeepEMhancer (26) was then used with the two unfiltered half maps to generate a locally sharpened map. These maps were then aligned with the full map and combined using the “vop maximum” function in UCSF Chimera based on the maximum value at each voxel (27) (Fig. S7). All resolutions were estimated by applying a soft mask around the protein density using the gold-standard Fourier shell correlation (FSC) = 0.143 criterion (28) (Fig. S4B).

### Cryo-EM model building, refinement, and analysis

Previously solved cryo-EM structures of human proteasomes were used for atomic model building (PDB 6MSE for the RP subcomplex of E_B_-like, PDB 7W38 for S_A_-like particles, and PDB 6MSK for E_D_-like particles) (8, 9, 19). For CP subcomplex, PDB 8CVR (20S, MG-132) was used as an initial model to build into the maps. Refined MIDN-26S proteasome model was used for building and refinement into the MIDN-26S proteasome without MG-132 map. All models were initially fitted individually as a rigid body model into each of the cryo-EM maps followed by adjustments of main-chains in Coot (29). Initial model for MIDN αHelix-C, Catch domain, and UBL domain was first derived from AlphaFold (30). The atomic models for RPN1-MIDN αHelix-C and RPN11-MIDN UBL domain were first rebuilt and refined against the high resolution local maps and again finally refined against the map of state E_B_^MIDN^. Catch domain was manually docked into low-pass filtered cryo-EM map (6 Å). Although we added MIDN-IRF4 complex into our sample we did not observe definitive density for IRF4 in our cryo-EM reconstructions; we did observe electron density in the AAA channel, which we modelled as a poly-peptide chain. Nucleotide densities were of sufficient quality for differentiating ADP from ATP in our S_A_^Apo^, E_B_^MIDN^ and E_D_^MIDN^ state maps, and allowed us to build the atomic models of ADP and ATP into their densities in the all the states.

Structure refinement was carried out in real space with PHENIX.real_space_refine (31) with global minimization applied with non-crystallographic symmetry (NCS), rotamer and Ramachandran constraints. Figures were generated with UCSF Chimera X (32). Interface area calculation was done using the PISA server (https://www.ebi.ac.uk/pdbe/pisa/). Refinement and model-building statistics are summarized in Table S2.

### Multiple sequence alignment

Multiple sequence alignment was performed using JalView (33).

## Data availability

The cryo-EM density maps and models generated in this study have been deposited in the EMDB and PDB databases under accession codes EMDB-49507/PDB-9NKF (S_A_^Apo^ composite map), EMDB-49508/PDB-9NKG (E_B_^MIDN^ composite map), EMDB-49509/PDB-9NKI (E_B_^MIDN-UBL^ composite map), EMDB-49510/PDB-9NKJ (E_D_^MIDN^ composite map). Locally refined maps and consensus maps of individuals states were deposited under EMDB-49497, EMDB-49498, EMDB-49498, EMDB-49499, EMDB-49500, EMDB-49501, EMDB-49502, EMDB-49503. EMDB-49504, EMDB-49505, and EMDB-49506. Plasmid information, cryo-EM micrographs, 2D classes, FSC curves, ResMap plots, and data processing 3D-classification and particle sorting schemes generated in this study are provided in the Supplementary Information.

## Acknowledgments

The authors would like to thank the UT Southwestern core facilities that have supported this work: Daniel Stoddard and the staff at the UT Southwestern Cryo-Electron Microscopy Facility, funded in part by the CPRIT Core Facility Support Award RP170644, and James Chen, Yan Han, Yang Li, and Joyce Fung at the UT Southwestern Structural Biology Lab. We thank Jan Erzberger (UT Southwestern) for valuable suggestions during data processing and analysis. This work was funded by National Institutes of Health grant R01AI125581 (BB).

---

**Figure S1.**
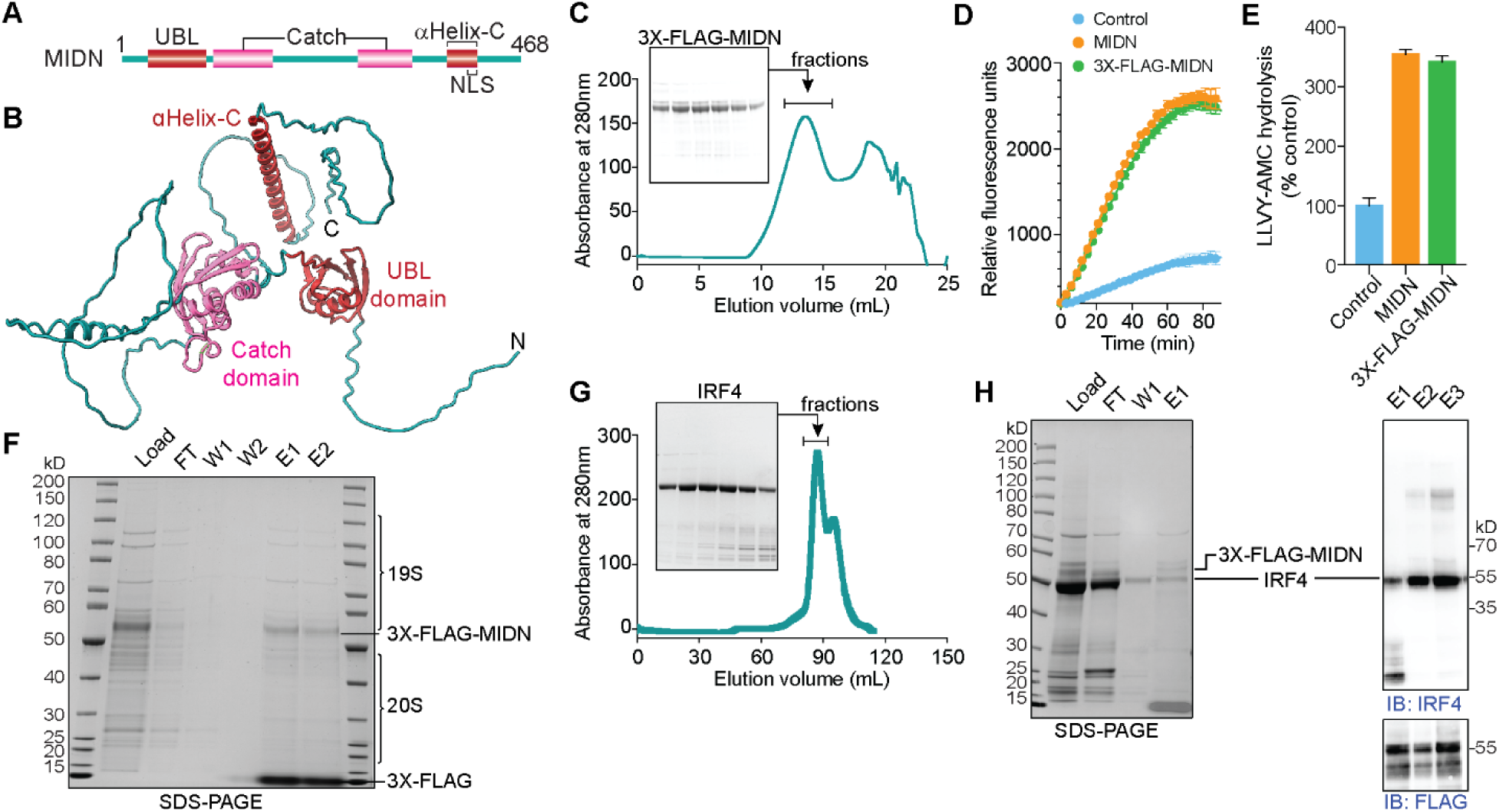
Protein purification and in vitro reconstitution of MIDN-proteasome complex. (A) Schematic representation of domain organization of MIDN (1-468 aa). UBL and αHelix-C (brick red), Catch domain (pink). (B) AlphaFold-predicted structure of MIDN (AF-Q504T8-F1), domains are colored according to schematic representation. (C) Size exclusion chromatography profile of MIDN purification on S200 column and inset shows SDS-PAGE analysis of peak fractions of MIDN protein. D) Time course analysis of proteasome peptidase activity towards the substrate LLVY-AMC in the absence (Control) or presence of MIDN or 3X-Flag-MIDN^WT^ as measured by AMC fluorescence. (E) Proteasome peptidase activity 80 min after initiation of the assay, plotted relative to the activity of proteasomes without added MIDN proteins (Control, set at 100%) (n = 3 reactions per condition). (F) SDS-PAGE analysis of pulldown using anti-Flag affinity gel of 26S proteasomes with 3X-Flag-MIDN^WT^. (G) Size exclusion chromatography profile of IRF4 purification on S200 column and inset shows SDS-PAGE analysis of peak fractions of IRF4 protein. (H) SDS-PAGE analysis (left) and immunoblot analysis (right) of pulldown using anti-Flag affinity gel of IRF4 with 3X-Flag-MIDN^WT^. FT, flow through; W, wash; E, elution.

**Figure S2.**
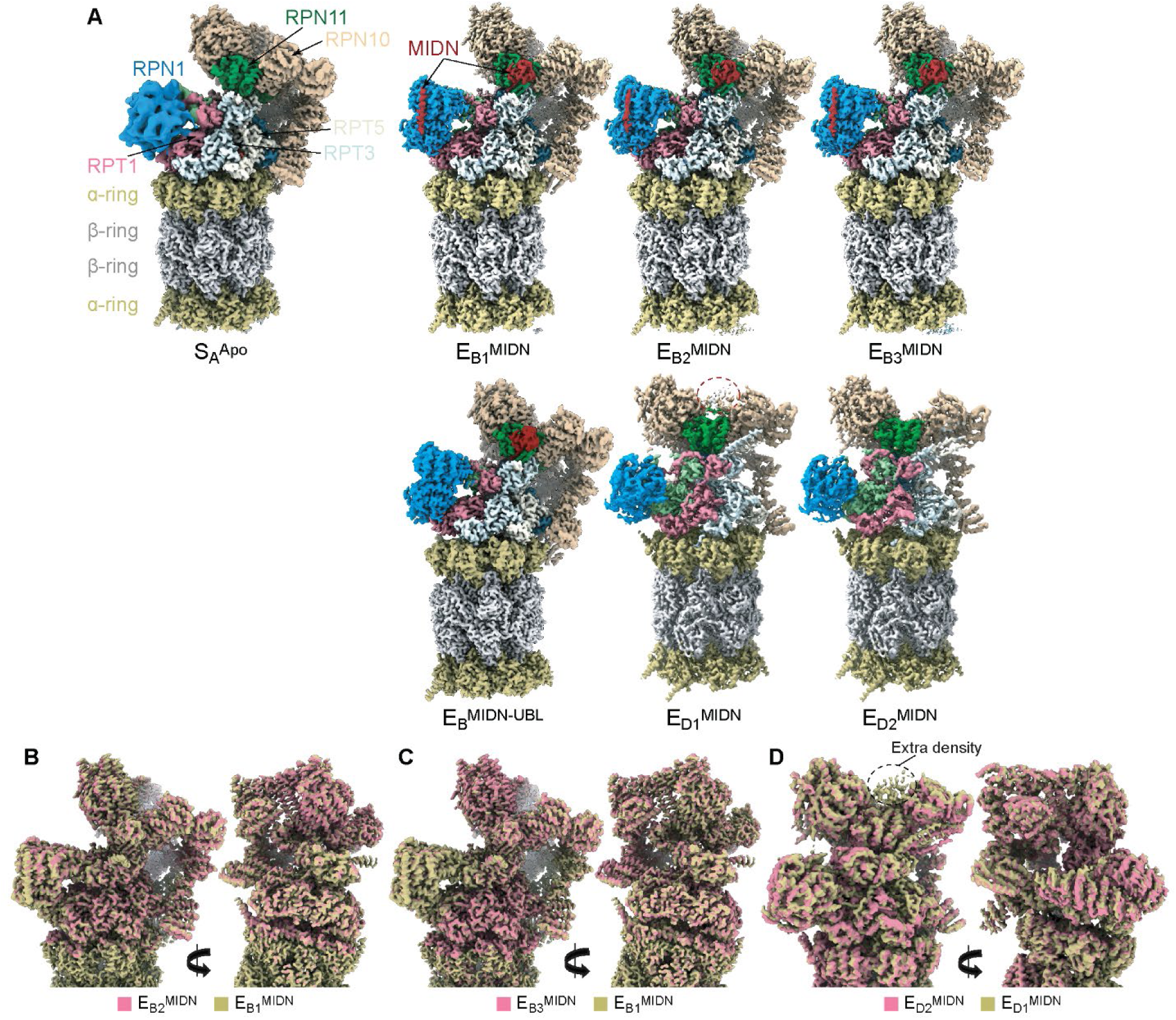
Conformational states of MIDN-bound and MIDN-free proteasomes. (A) Cryo-EM maps of MIDN- and substrate-free 26S proteasome S_A_-like state (designated S_A_^Apo^), MIDN-bound E_B_-like states (designated E_B1_^MIDN^, E_B2_^MIDN^, E_B3_^MIDN^, and E_B_^MIDN-UBL^), and MIDN-bound E_D_-like proteasome states (designated E_D1_^MIDN^ and E_D2_^MIDN^). Extra density in E_D1_ state is shown in a dashed circle. (B-D) Comparison of cryo-EM reconstructions. (B) E_B1_^MIDN^ with E_B2_^MIDN^. (C) E_B1_^MIDN^ with E_B3_^MIDN^. (D) E_D1_^MIDN^ with E_D2_^MIDN^.

**Figure S3.**
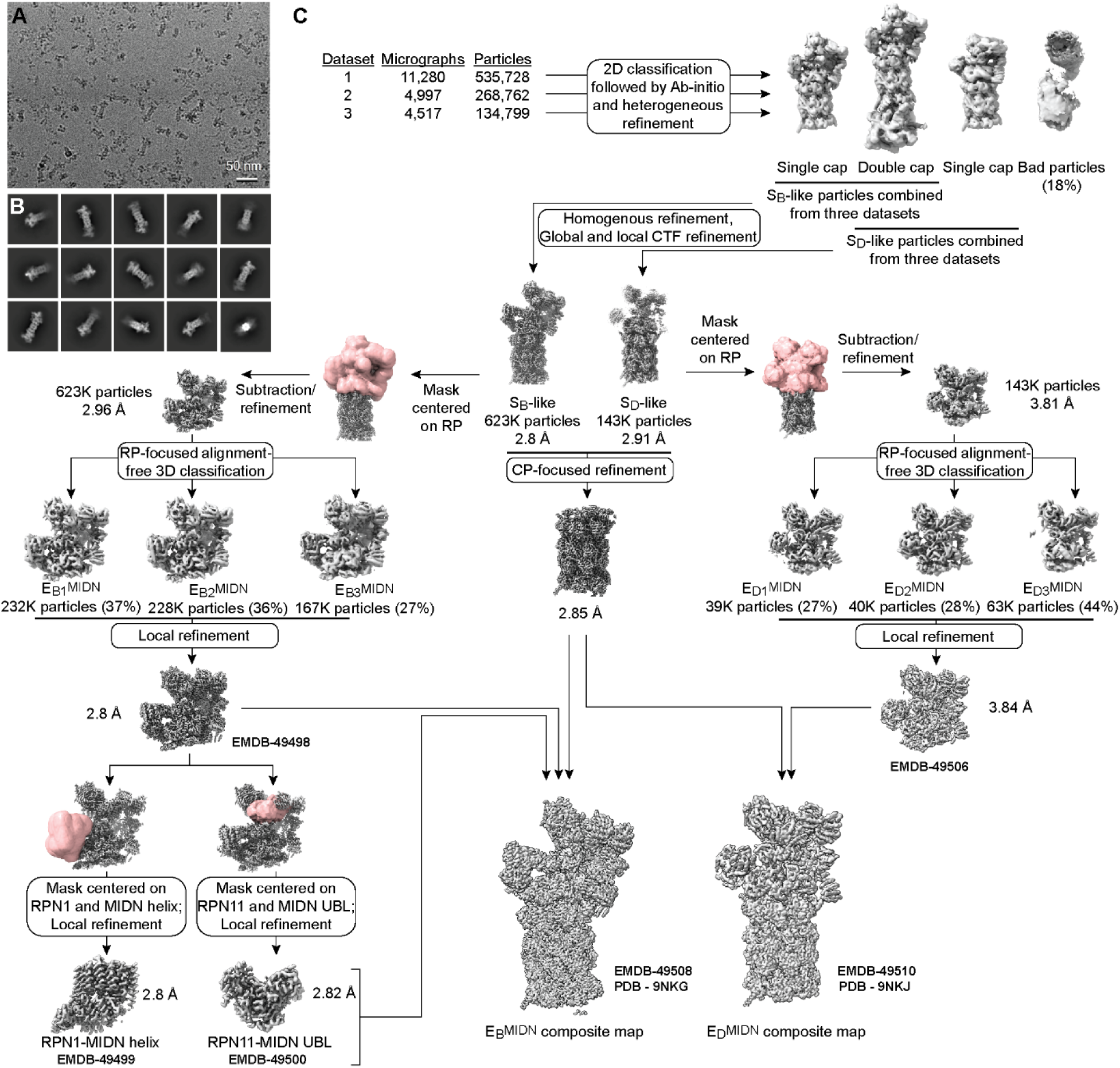
Cryo-EM imaging and data processing workflow for MIDN-IRF4-proteasome complex in the presence of ATP*γ*S and MG-132. (A) Representative cryo-EM micrograph out of a total 20,794 collected. Data from three separate grids frozen at the same time (identical sample) were combined. (B) Representative unsupervised 2D classes with a broad distribution of orientations showing clear secondary structure features were selected for further processing. (C) Workflow diagram of 3D classification strategy and the final composite maps of the MIDN-bound 26S proteasome in E_B_^MIDN^ and E_D_^MIDN^ states. Map volumes are shown for all steps and masks (red) indicate volume selected for local classification. Major sorting and classification criteria, particle numbers and percentages (for each 3D classification), and final map resolutions are indicated. The accession numbers are included for maps deposited to PDB/EMDB.

**Figure S4.**
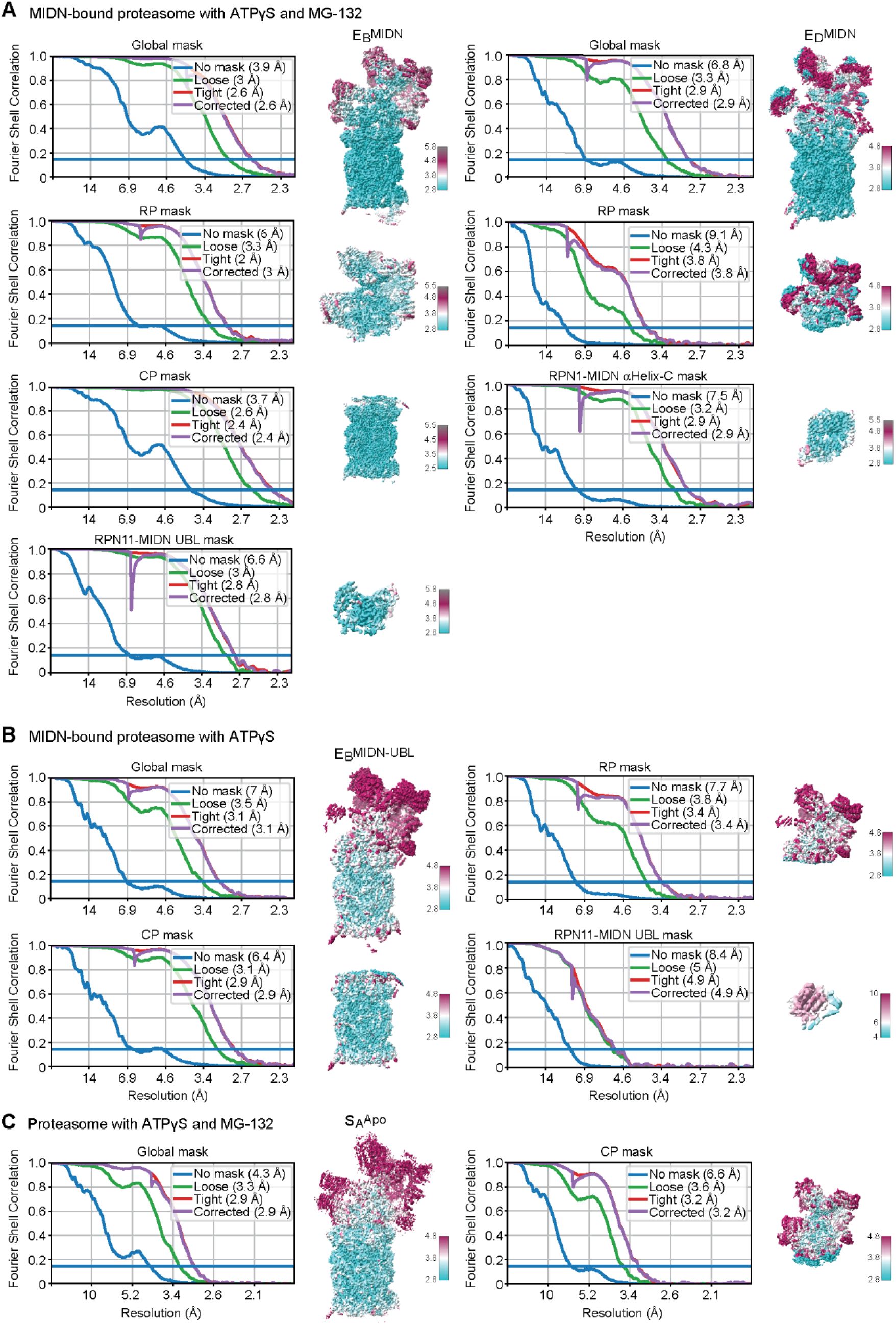
Fourier shell correlation (FSC) curves and local resolution plots. Gold standard FSC plots and corresponding local resolution estimates for 26S, 19S, and 20S proteasome complexes, and complexes of RPN1-MIDN αHelix-C and RPN11-MIDN UBL domain in (A) E_B_^MIDN^ state (left) and E_D_^MIDN^ state (right), (B) E_B_^MIDN-UBL^ state, and (C) S_A_^Apo^ state.

**Figure S5.**
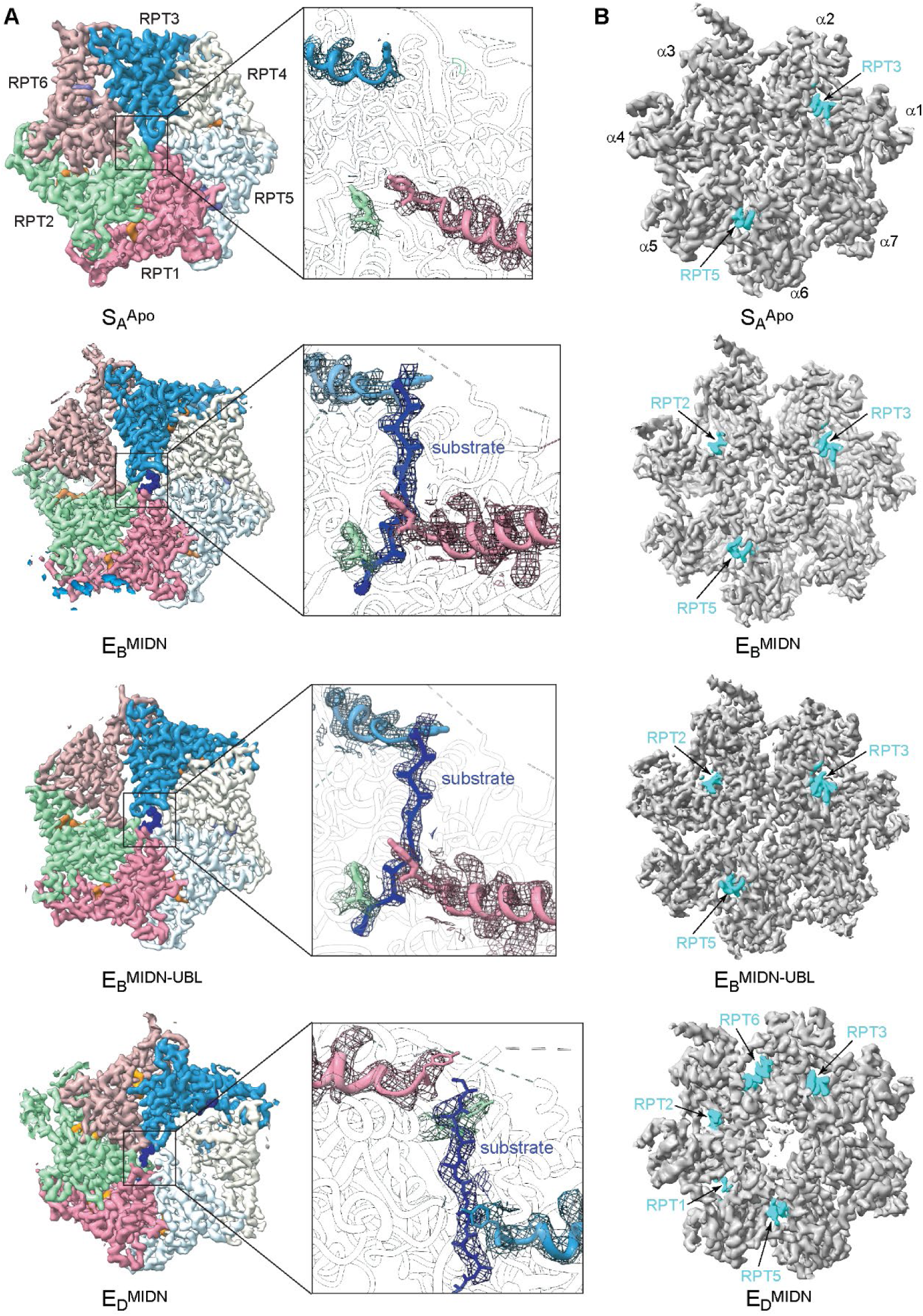
Substrate interactions with the pore loops in AAA ATPase motor and the RP-CP interface. (A) Cryo-EM density of the ATPase ring in different proteasome states shown as surfaces and density of the substrate polypeptide backbone represented as blue sticks (left). Close-up view of the substrate density shown as a mesh (right). (B) Overview of the RP–CP interface in which the carboxyl-terminal tails of RPT subunits are inserted into the α-pockets of the CP in different states. The cryo-EM density of the CP subcomplex is shown as a grey surface and RPT C-tail densities are represented as teal blue surfaces.

**Figure S6.**
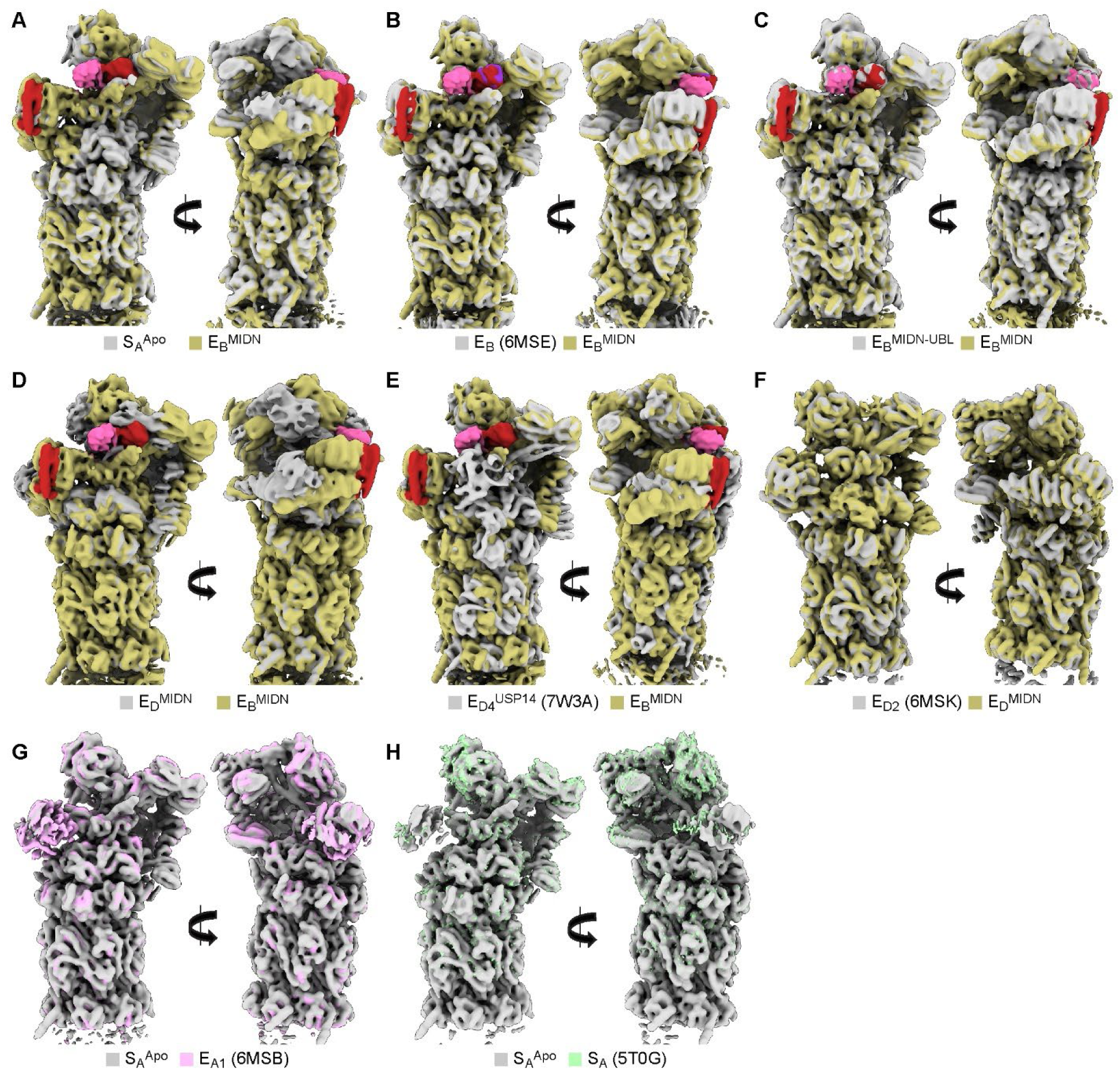
Structural comparison of MIDN-bound and MIDN-free 26S proteasomes with each other and with published proteasome states. (A) S_A_^Apo^ state (grey) with E_B_^MIDN^ state (dark khaki), (B) E_B_^MIDN^ state (dark khaki) with substrate-engaged MIDN-free E_B_ state (6MSE), (C) substrate-engaged MIDN-bound E_B_ state in the presence of ATPγS and MG-132 (E_B_^MIDN^, dark khaki) and the same without MG-132 (E_B_^MIDN-UBL^, grey). (D) E_B_^MIDN^ state (dark khaki) with E_D_^MIDN^ (grey), (E) E_B_^MIDN^ state (dark khaki) with USP14-bound proteasome structure E_D4_^USP14^ (7W3A) (grey), (F) E_D_^MIDN^ state (dark khaki) with processive substrate translocation E_D2_ proteasome state (6MSK) (grey), (G) S_A_^Apo^ state (grey) with the substrate-bound E_A1_ state (6MSB) (pink), and (H) S_A_^Apo^ state (grey) with the substrate-free S_A_ state of the human proteasome (5T0G) (green).

**Figure S7.**
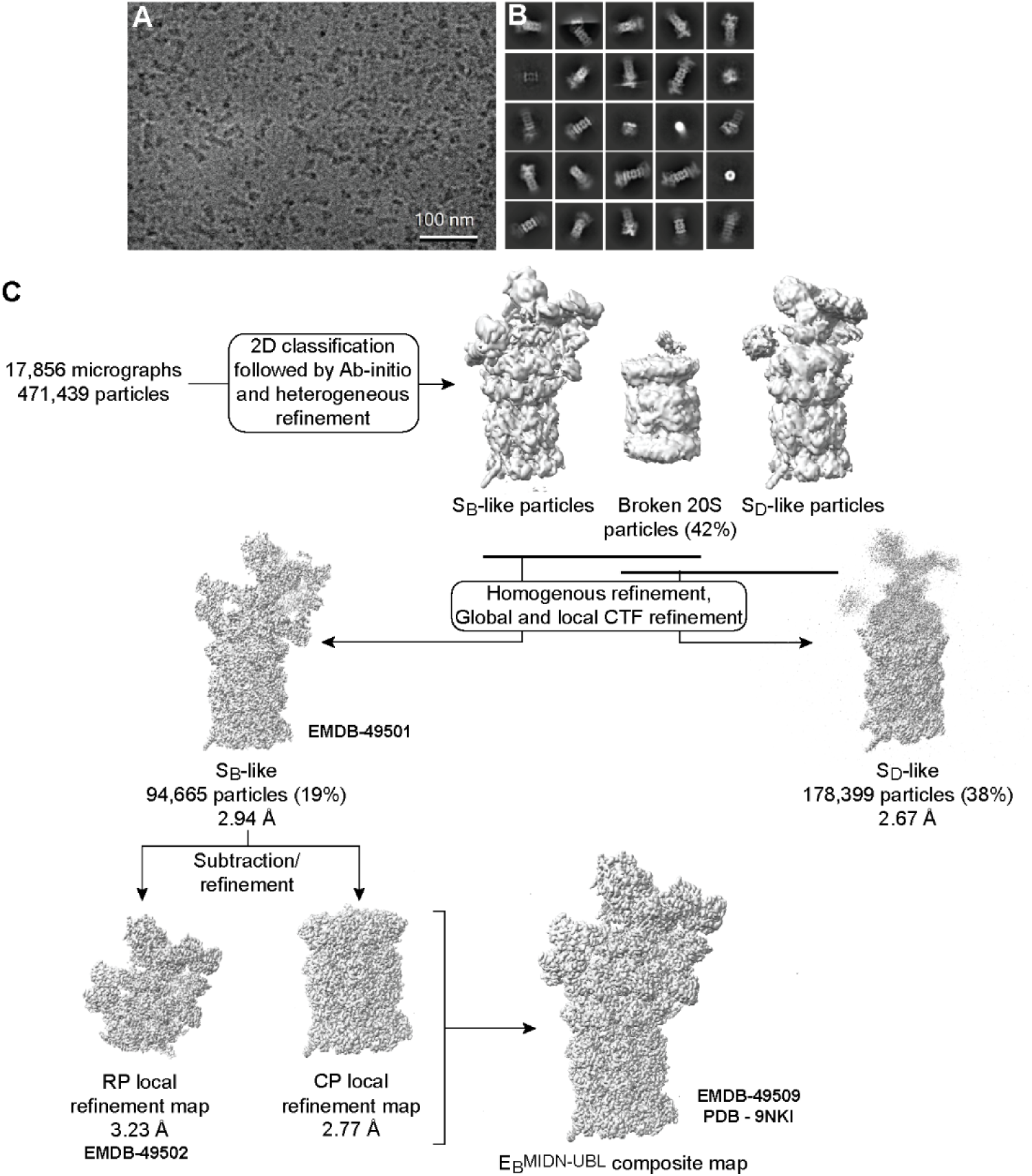
Cryo-EM imaging and data processing workflow for MIDN-IRF4-proteasome complex in the presence of ATP*γ*S. (A) Representative cryo-EM micrograph out of 17,856 images collected. (B) Representative unsupervised 2D classes with a broad distribution of orientations showing clear secondary structure features were selected for further processing. (C) Workflow diagram of 3D classification strategy and the final composite maps of the MIDN-bound 26S proteasome in E_B_^MIDN-UBL^ and S_D_-like states. Map volumes are shown for all steps. Major sorting and classification criteria, particle numbers and percentages (for each 3D classification), and final map resolutions are indicated. The accession numbers are included for maps deposited to PDB/EMDB.

**Figure S8.**
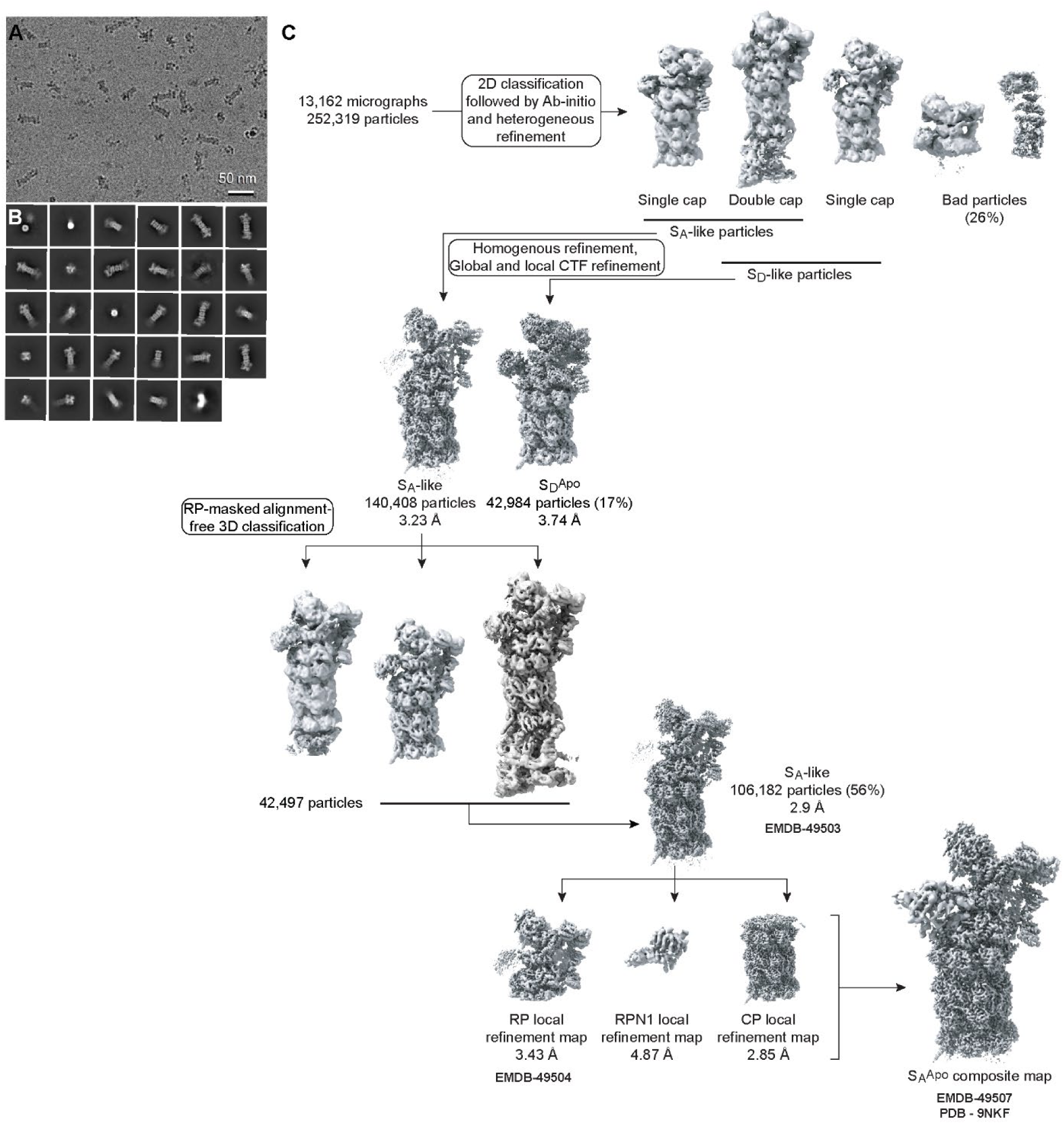
Cryo-EM imaging and data processing workflow for MIDN- and substrate-free 26S proteasome complex in the presence of ATP*γ*S and MG-132. (A) Representative cryo-EM micrograph out of 13,162 images collected. (B) Representative unsupervised 2D classes with a broad distribution of orientations showing clear secondary structure features were selected for further processing. (C) Workflow diagram of 3D classification strategy and the final composite maps of the MIDN-free 26S proteasome in S_A_^Apo^ state. Map volumes are shown for all steps. Major sorting and classification criteria, particle numbers and percentages (for each 3D classification), and final map resolutions are indicated. The accession numbers are included for maps deposited to PDB/EMDB.

**Figure S9.**
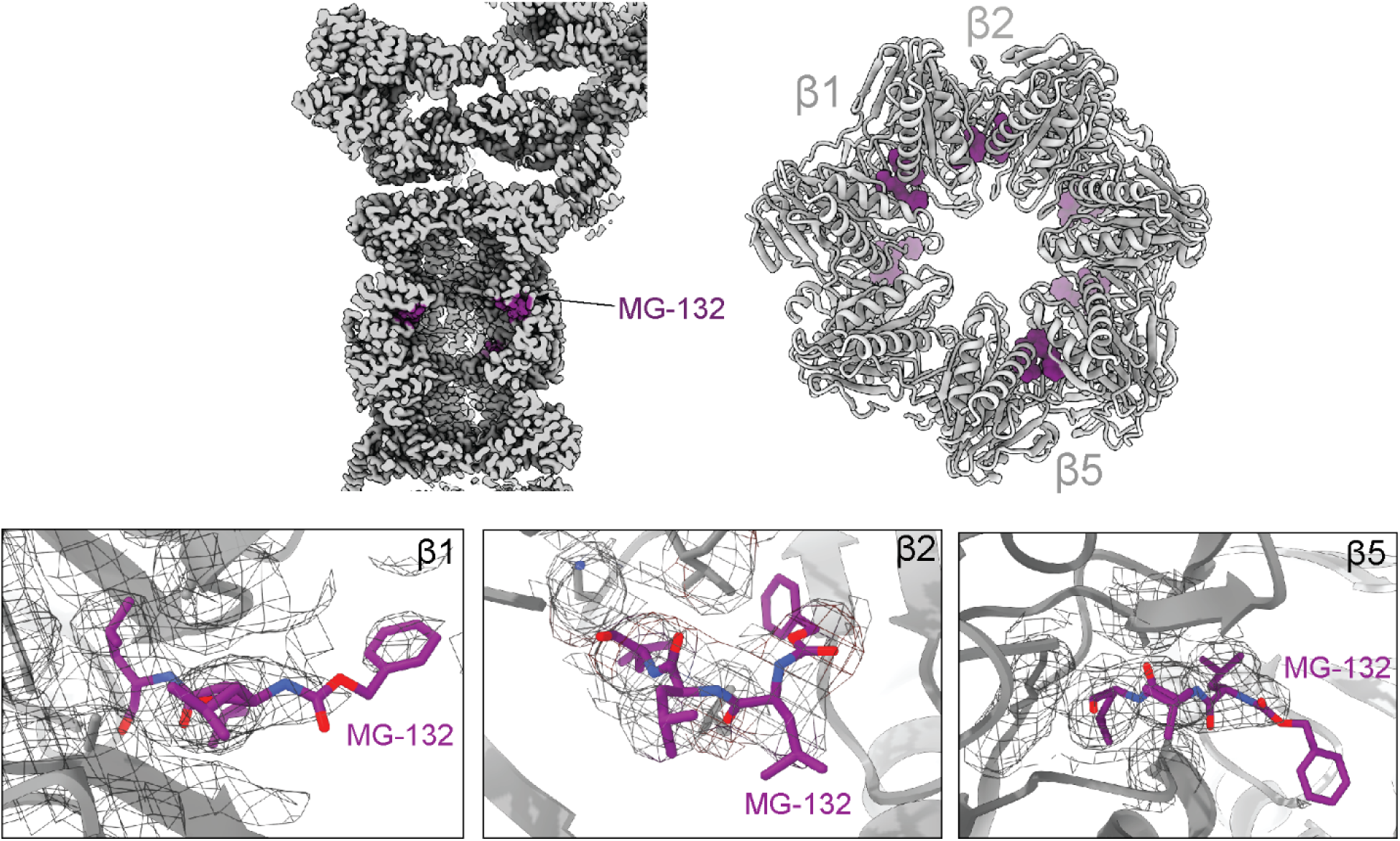
MG-132 cryo-EM density. MG-132 (purple) bound to three β-subunits of the CP (grey) in E_B_^MIDN^ state (upper). Cryo-EM densities are shown in grey mesh (lower).

**Figure S10.**
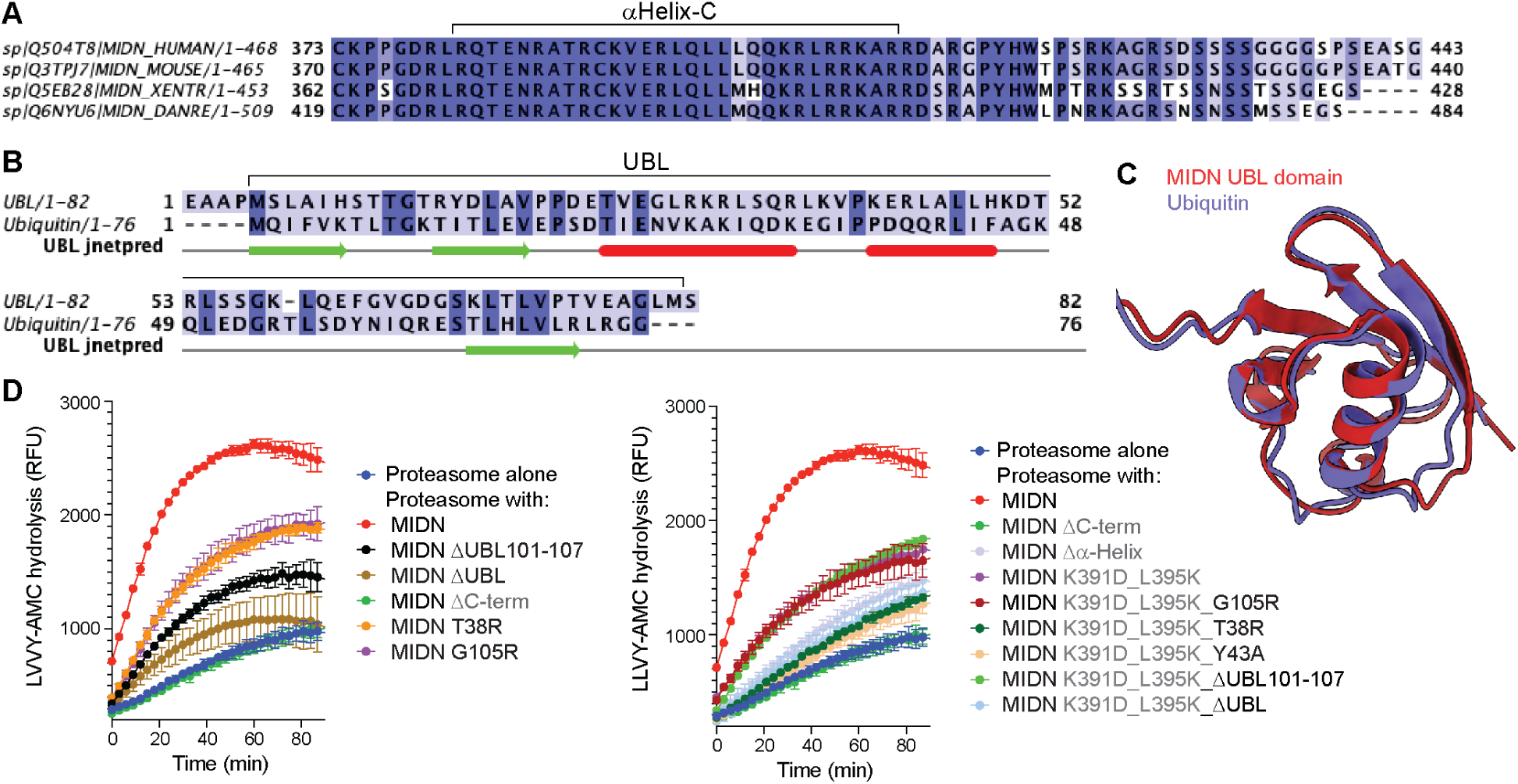
Conservation of MIDN *α*Helix-C amino acid sequences across species and structural similarity of the MIDN UBL domain and ubiquitin. (A) Multiple sequence alignment of MIDN αHelix-C amino acid sequences from different species. (B) Amino acid sequence alignment of MIDN-UBL domain with ubiquitin. Predicted secondary structure is shown below (green, β-strand; red, α-helix). Residues are colored based upon conservation score (dark, highly conserved; light, poorly conserved) (B and C). (C) Overlay of MIDN-UBL domain structure (brick red) with ubiquitin structure (purple) (1UBQ) shows that they adopt a similar β-grasp fold. (D) Time course analysis of proteasome peptidase activity towards the substrate LLVY-AMC in the absence (Proteasome alone) or presence of WT MIDN or MIDN mutant proteins as measured by AMC fluorescence (n = 3 reactions per condition). MIDN UBL domain mutations are labelled in black; MIDN αHelix-C mutations are labelled in grey.

**Figure S11.**
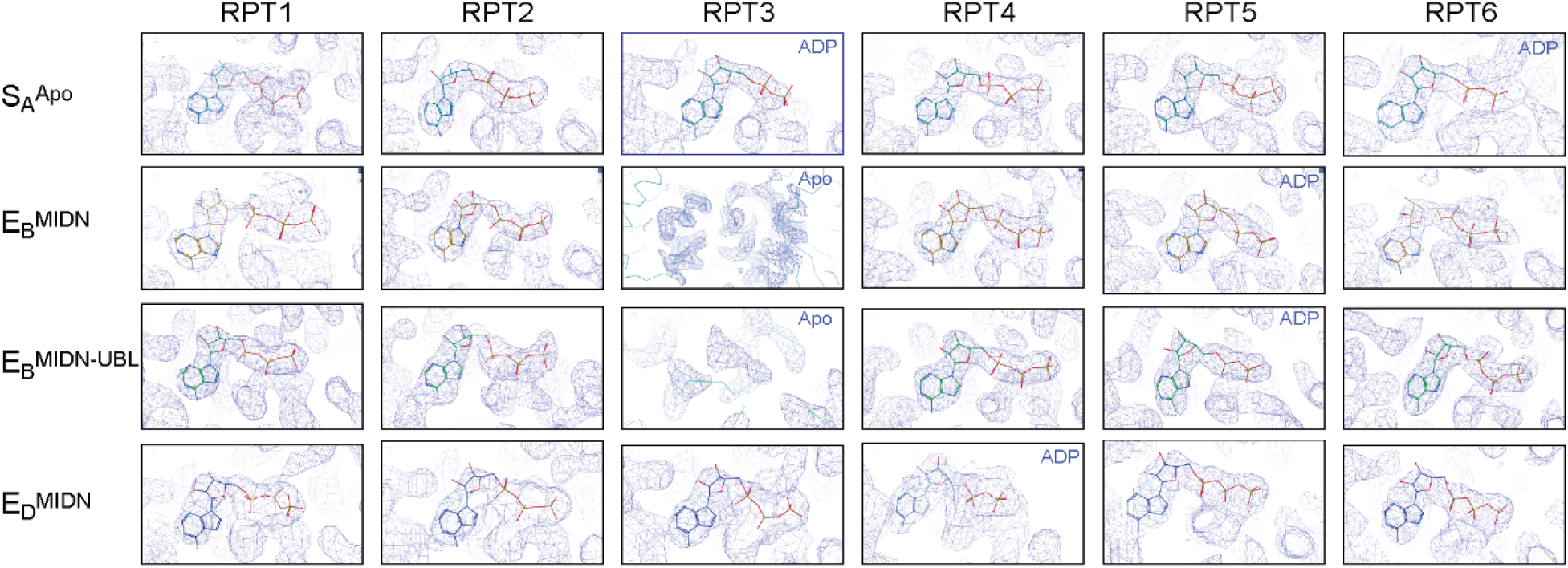
Nucleotide densities in the ATPases of the RP in different proteasome states. Comparison of the nucleotide densities in the cryo-EM reconstructions of MIDN-free and MIDN-bound proteasomes. The nucleotide densities are shown in blue mesh fitted with atomic models. All views are screenshots directly from coot (ref. 29). ADP-bound and Apo states are labelled; all others are ATP-bound.

**Table S1.**
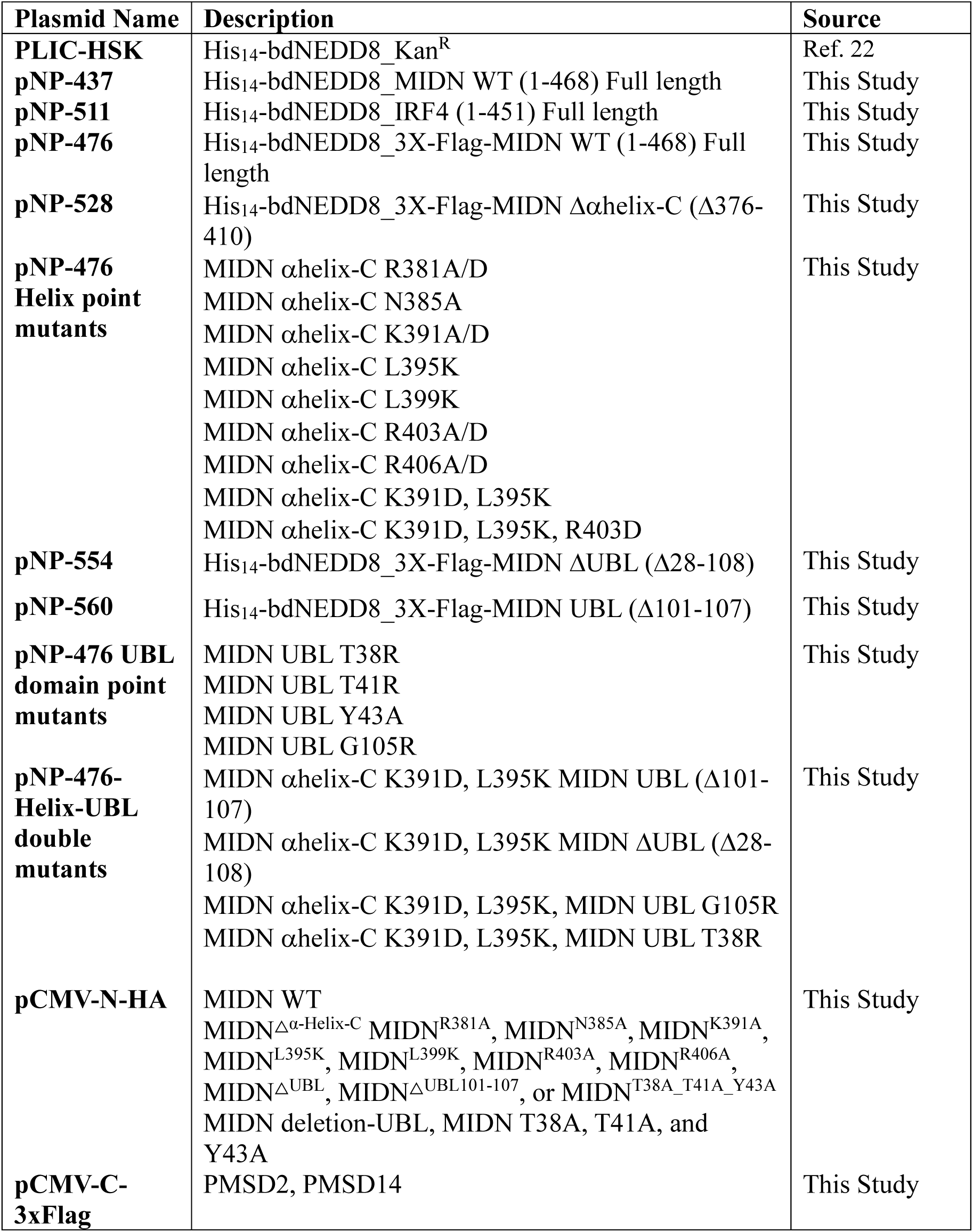
Plasmids.

**Table S2.** Cryo-EM data collection, refinement, and validation statistics.

|  | **S<sub>A</sub><sup>Apo</sup>** | **E<sub>B</sub><sup>MIDN</sup>** | **E<sub>B</sub><sup>MIDN_UBL</sup>** | **E<sub>D</sub><sup>MIDN</sup>** |
| --- | --- | --- | --- | --- |
|  | PDB:9NKF | PDB:9NKG | PDB: 9NKI | PDB:9NKJ |
|  | EMDB-49507 | EMDB-49508 | EMDB-49509 | EMDB-49510 |
| **Data collection and Processing** |  |  |  |  |
| Voltage (keV) | 300 | 300 | 300 | 300 |
| Camera | Gatan | Gatan | Falcon (4i) | Gatan |
| Magnification | 81000x | 81000x | 105000x | 81000x |
| Pixel size at detector (Å/pixel) | 1.08 | 1.08 | 0.73 | 1.08 |
| Total electron exposure (e<sup>-</sup>/Å<sup>2</sup>) | 50 | 50 | 50 | 50 |
| Exposure rate (e<sup>-</sup>/pixel/sec) | 8 | 8 | 7.76 | 8 |
| Number of frames collected | 44 | 44 | 1092 | 44 |
| Defocus range (μm) | -1.1 to -2.7 | -1.1 to -2.7 | -1.1 to -2.7 | -1.1 to -2.7 |
| Automation software | SerialEM | SerialEM | SerialEM | SerialEM |
| Energy filter slit width | 10 | 10 | 10 | 10 |
| Micrographs collected (no.) | 13,162 | 20,794 | 17856 | 20,794 |
| Total extracted particles (no.) | 252,313 | 939,289 | 471,439 | 939,289 |
| Refined particles (no.) | 142,000 | 623,000 | 272,000 | 142,000 |
| Final particles (no.) | 106,000 | 232,000 | 94,000 | 39,000 |
| Symmetry imposed | C1 | C1 | C1 | C1 |
| Resolution (global, Å) | 2.9 | 2.9 | 2.8 | 3.8 |
| FSC threshold | 0.143 | 0.143 | 0.143 | 0.143 |
| **Refinement** |  |  |  |  |
| Refinement package | Phenix | Phenix | Phenix | Phenix |
| Initial Model used | 7W38 | 6MSE | 6MSE | 6MSK |
| Model resolution (Å) | 3.2 | 2.9 | 3.2 | 4.1 |
| FSC Threshold | 0.5 | 0.5 | 0.5 | 0.5 |
| **Model composition** |  |  |  |  |
| Non-hydrogen atoms | 101342 | 103530 | 102622 | 102,224 |
| Protein | 13233 | 13430 | 13398 | 13321 |
| Ligands | 18 | 16 | 10 | 15 |
| **B factors (Å<sup>2</sup>)** |  |  |  |  |
| Protein residues | 66 | 66 | 73 | 82.7 |
| Ligands | 53 | 60 | 46 | 63 |
| **R.m.s. deviations** |  |  |  |  |
| Bond lengths (Å) | 0.004 | 0.003 | 0.003 | 0.004 |
| Bond angles (°) | 0.545 | 0.528 | 0.573 | 0.59 |
| **Validation** |  |  |  |  |
| MolProbity score | 2.86 | 1.91 | 1.96 | 3.08 |
| CaBLAM outliers | 3.66 | 2.52 | 2.55 | 5.15 |
| Clashscore | 33.26 | 6.97 | 9.88 | 39.8 |
| Poor rotamers (%) | 3.67 | 1.95 | 1.63 | 3.89 |
| **Ramachandran plot** |  |  |  |  |
| Favored (%) | 93.6 | 95.5 | 95.8 | 90 |
| Allowed (%) | 6.2 | 4.33 | 4 | 9.55 |
| Outliers (%) | 0.18 | 0.12 | 0.11 | 0.45 |

